# Fast signaling and focal connectivity of PV^+^ interneurons ensure efficient pattern separation by lateral inhibition in a full-scale dentate gyrus network model

**DOI:** 10.1101/647800

**Authors:** Segundo Jose Guzman, Alois Schlögl, Claudia Espinoza, Xiaomin Zhang, Ben Suter, Peter Jonas

**Affiliations:** IST Austria (Institute of Science and Technology Austria), Am Campus 1, A-3400 Klosterneuburg, Austria; Institute for Molecular Biotechnology (IMBA), Dr. Bohr-Gasse 3, A-1030 Wien, Austria

**Keywords:** GABAergic interneurons, PV^+^ interneurons, lateral inhibition, granule cells, hippocampus, dentate gyrus, pattern separation, winner-takes-all mechanism

## Abstract

Pattern separation is a fundamental brain computation that converts small differences in synaptic input patterns into large differences in action potential (AP) output patterns. Pattern separation plays a key role in the dentate gyrus, enabling the efficient storage and recall of memories in downstream hippocampal CA3 networks. Several mechanisms for pattern separation have been proposed, including expansion of coding space, sparsification of neuronal activity, and simple thresholding mechanisms. Alternatively, a winner-takes-all mechanism, in which the most excited cells inhibit all less-excited cells by lateral inhibition, might be involved. Although such a mechanism is computationally powerful, it remains unclear whether it operates in biological networks. Here, we develop a full-scale network model of the dentate gyrus, comprised of granule cells (GCs) and parvalbumin^+^ (PV^+^) inhibitory interneurons, based on experimentally determined biophysical cellular properties and synaptic connectivity rules. Our results demonstrate that a biologically realistic principal neuron–interneuron (PN–IN) network model is a highly efficient pattern separator. Mechanistic dissection in the model revealed that a winner-takes-all mechanism by lateral inhibition plays a crucial role in pattern separation. Furthermore, both fast signaling properties of PV^+^ interneurons and focal GC–interneuron connectivity are essential for efficient pattern separation. Thus, PV^+^ interneurons are not only involved in basic microcircuit functions, but also contribute to higher-order computations in neuronal networks, such as pattern separation.

## INTRODUCTION

A fundamental question in neuroscience is to understand how higher-order computations in the brain are implemented at the level of synapses, neurons, and neuronal networks. A key computation in the brain is pattern separation, a process that converts slightly different input patterns into highly different action potential (AP) output patterns^1–3^. Pattern separation is thought to play a particularly important role in the memory circuits of the hippocampus, where separation computations at the input layer, the dentate gyrus^4^, facilitate reliable storage and recall of memories in the downstream layer, the CA3 region^2, 5–7^. However, although pattern separation has an important function in memory circuits, the underlying mechanisms remain elusive.

In the cerebellum, a circuit where pattern separation is relevant for precise motor control^8^, synaptic divergence from a small to a large number of neurons and sparsification of activity are key factors^9–11^. However, as the connectivity between synaptic input and cerebellar granule cells (GCs) is extremely sparse^11^, generalization to the dentate gyrus is not straightforward. In the olfactory bulb, a circuit where pattern separation converts broad activation of sensory olfactory neurons into specific activation of mitral cells, a winner-takes-all mechanism mediated by lateral inhibition contributes to pattern separation^12–21^. However, in olfactory circuits lateral inhibition is mediated by specialized dendro-dendritic synapses, and the number of inhibitory GCs exceeds the number of excitatory mitral cells by more than an order of magnitude^22^. Whether lateral inhibition contributes to pattern separation in the dentate gyrus, where signaling is mediated by axo-dendritic synapses and excitatory neurons greatly outnumber inhibitory cells^23^, remains unclear^24^.

We recently found that in the dentate gyrus lateral inhibition by parvalbumin-expressing (PV^+^) interneurons is more abundant than in any other studied brain region^25^, consistent with the idea that lateral inhibition implements a winner-takes-all mechanism underlying pattern separation^25^. However, principal neuron–interneuron (PN–IN) connectivity in the dentate gyrus is highly focal, which seems incompatible with the central idea of that model, that a winner should be able to globally suppress all non-winners. To clarify the role of lateral inhibition in pattern separation in the dentate gyrus, we constructed a network model of this brain area based on experimentally determined biophysical cellular properties and synaptic connectivity rules. In contrast to several previous studies, the model was implemented in full-scale. We quantitatively analyzed pattern separation in the model to address three main questions. First, is a PN–IN network with biological properties able to perform efficient pattern separation? Second, what is the role of lateral inhibition in pattern separation? Third, how do the fast signaling properties of GABAergic interneurons^26^ and the focal PN–IN connectivity^25^ impact on pattern separation? A preliminary account of this work has been published in abstract form^27^.

## RESULTS

### A winner takes-all-mechanism is able to decorrelate patterns

Pattern separation is a network computation that converts highly overlapping synaptic input patterns into minimally overlapping AP output patterns. The basic principle is illustrated in Fig. 1a. When two highly overlapping input patterns (A and B) are applied at the input of a neuronal population (Fig. 1a, top), two largely non-overlapping output patterns (A’ and B’) are generated at the output of the population (Fig. 1a, bottom). Quantitatively, the correlation coefficients for the output patterns (*R*_out_ = r(A’, B’)) are smaller than the corresponding correlation coefficients of the input patterns (*R*_in_ = r(A, B)). Thus, when *R*_out_ is plotted against *R*_in_ for all pairs of patterns, data points should be located below the identity line (Fig. 1b).

**Fig. 1.**
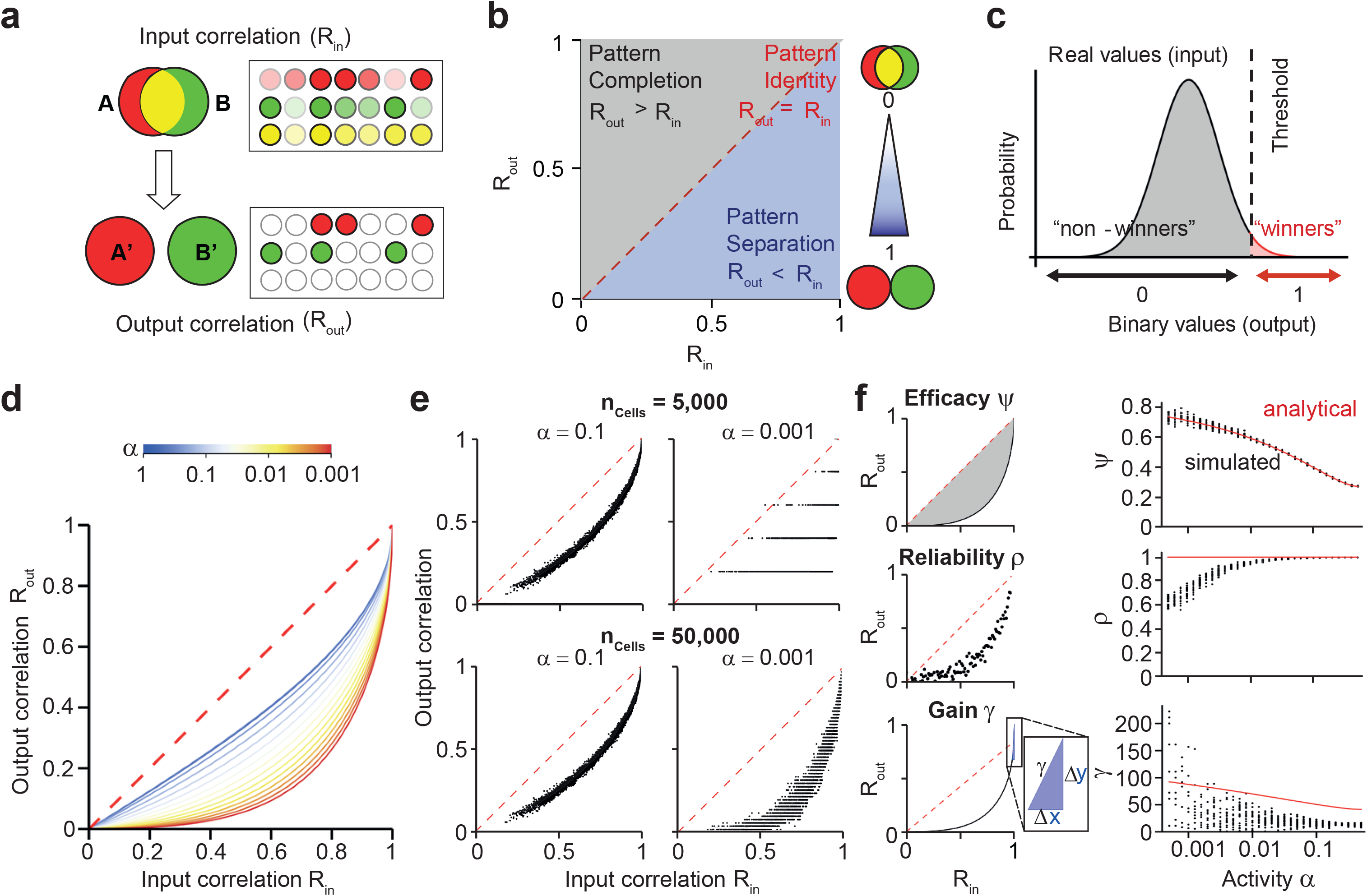
Principles and quantitative analysis of pattern separation by a winner-takes-all mechanism. (**a**) Left, Venn diagrams of two patterns before and after pattern separation. Overlapping input patterns (A, B; top) are converted into non-overlapping output patterns (A’, B’; bottom). Right, real input pattern vectors and binary output pattern vectors; red, cells active in pattern A; green, cells active in pattern B; yellow, cells active in both patterns. In top scheme, intensity reflects the level of excitatory synaptic drive. (**b**) Analysis of pattern separation in input-output correlation plots. *R*_in_ and *R*_out_ represent pairwise correlations in input and output patterns. Red dashed line indicates pattern identity. Area below identity line corresponds to a regime in which *R*_out_ < *R*_in_, i.e. pattern separation. Area above identity line corresponds to a regime where *R*_out_ > *R*_in_, i.e. pattern completion. (**c**) Simple representation of the winner-takes-all pattern separation mechanism. Real vectors corresponding to the excitatory synaptic drive (input pattern; Gaussian distribution) are converted into binary vectors representing the AP activity of the neurons (output pattern), applying a threshold to the data. The threshold is set to give a defined average activity level α. (**d**) *R*_out_ versus *R*_in_ plots for various activity levels α, varied in the range from 0.63 to 0.001. The normalized area between the curves and the identity line (pattern separation index 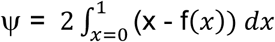 represents a robust measure of the efficacy of the pattern separation process. (**e**) *R*_out_ versus *R*_in_ plots for random vectors of finite neuronal populations with different numbers of cells *n*_Cells_. Top, *n*_Cells_ = 5,000; bottom, *n*_Cells_ = 50,000. Note that the correlation between *R*_out_ and *R*_in_ values drops for decreasing activity levels for *n*_Cells_ = 5,000, but remains stable for *n*_Cells_ = 50,000. This highlights the importance of full-scale simulations for understanding the mechanisms of pattern separation. (**f**) Three metrics to characterize pattern separation. Top, efficiency Ψ of pattern separation, quantified from the area under the *R*_out_–*R*_in_ curve. Center, reliability of pattern separation ρ, quantified by the Pearson’s correlation coefficient of the ranks of all *R*_out_ versus the ranks of all *R*_in_ values. Bottom, gain γ of pattern separation, determined from the slope of the *R*_out_–*R*_in_ curve for *R*_in_ → 1. Left, schematic illustration of definition of the parameters. Right, plot of Ψ, ρ, and γ against average activity α. Red curve, analytical solution for the infinite-size network; black points, 20 simulations per activity level, each for a population of 5,000 neurons. For the γ–α plot, only positive γ values are depicted.

To test these predictions, we used the simplest possible implementation of a winner-takes-all mechanism: an infinite-size network incorporating a thresholding mechanism (Fig. 1c). Under the assumption that input patterns follow a bivariate Gaussian distribution (Fig. 1c), *R*_out_ can be analytically computed for any given *R*_in_ and average activity level α using Hoeffding’s lemma^28^ (see Methods). As expected for a pattern separation mechanism, *R*_out_–*R*_in_ curves were consistently located below the identity line (Fig. 1d). To assess whether this mechanism also works in finite-size networks, we performed numerical simulations of input and output patterns (Fig. 1e). Random real number input patterns were drawn from a bivariate Gaussian distribution. Interestingly, the parameter dependence was more complex than predicted from the analytical solution for the infinite-size network. For small neuronal populations (*n*_Cells_= 5,000), reduction of activity α increased Ψ. However, below a certain activity level, the monotonic relation between *R*_out_ and *R*_in_ was disrupted (Fig. 1e, top, right). In contrast, for larger neuronal populations (*n*_Cells_ = 50,000), the monotonic relation between input and output was maintained over a wider range (Fig. 1e, bottom).

To characterize these complex phenomena, we introduced three quantitative measures of pattern separation (Fig. 1f; see Methods). First, we measured the efficacy of pattern separation Ψ as the normalized area between the data points and the identity line (Fig. 1f, top). Second, we computed the reliability of pattern separation ρ from the rank correlation coefficient of the *R*_out_–*R*_in_ data (Fig. 1f, center). Finally, we determined the maximal gain of pattern separation γ from the slope of the input-output correlation for *R*_in_ → 1 (Fig. 1f, bottom). For the infinite-size networks, Ψ approached values as high as 0.75 for low values of α. For the finite-size networks, pattern separation efficacy Ψ approached similar values. However, pattern separation reliability ρ was markedly reduced for low levels of α (ρ = 0.74 for α = 0.001 and *n*_Cells_ = 5,000; ρ = 0.94 for α = 0.001 and *n*_Cells_ = 50,000). In conclusion, these results provide a proof-of-principle that a winner-takes-all mechanism is able to separate patterns. However, the performance of the mechanism depends on activity level and network size.

### A biologically realistic PN–IN network model is an efficient pattern separator

To explore whether the winner-takes-all mechanism of pattern separation works in biologically realistic networks resembling the dentate gyrus, we developed a model of pattern separation based on empirical experimental data (Fig. 2; Supplementary Figure 1; Table 1). The network was created in full-scale, with 500,000 GCs^29^, represented as leaky integrate-and-fire neurons, and 2,500 PV^+^ interneurons, implemented as single-compartment conductance-based models (Fig. 2a). Excitatory GC–PV^+^ interneuron synapses, inhibitory PV^+^ interneuron–GC synapses, mutual inhibition, and gap junctions were implemented based on the detailed description of functional connectivity obtained by multi-cell recordings^25^. At the network input, 50,000 entorhinal cortex cells (ECs) were attached^23^. The EC–GC connectivity was constrained by the width of the entorhinal cortex neuron axons (20% of the dentate gyrus along the longitudinal axis)^30^ and the number of spines on the dendrites of GCs (~5,000)^31,32^. As gamma oscillations may contribute to a winner-takes-all mechanism^17^, an inhibitory conductance was initiated at the onset of each simulation epoch^17,33^. Since gamma oscillations show high power in the dentate gyrus^34–36^, this also contributed to the realism of the model.

**Table 1.**
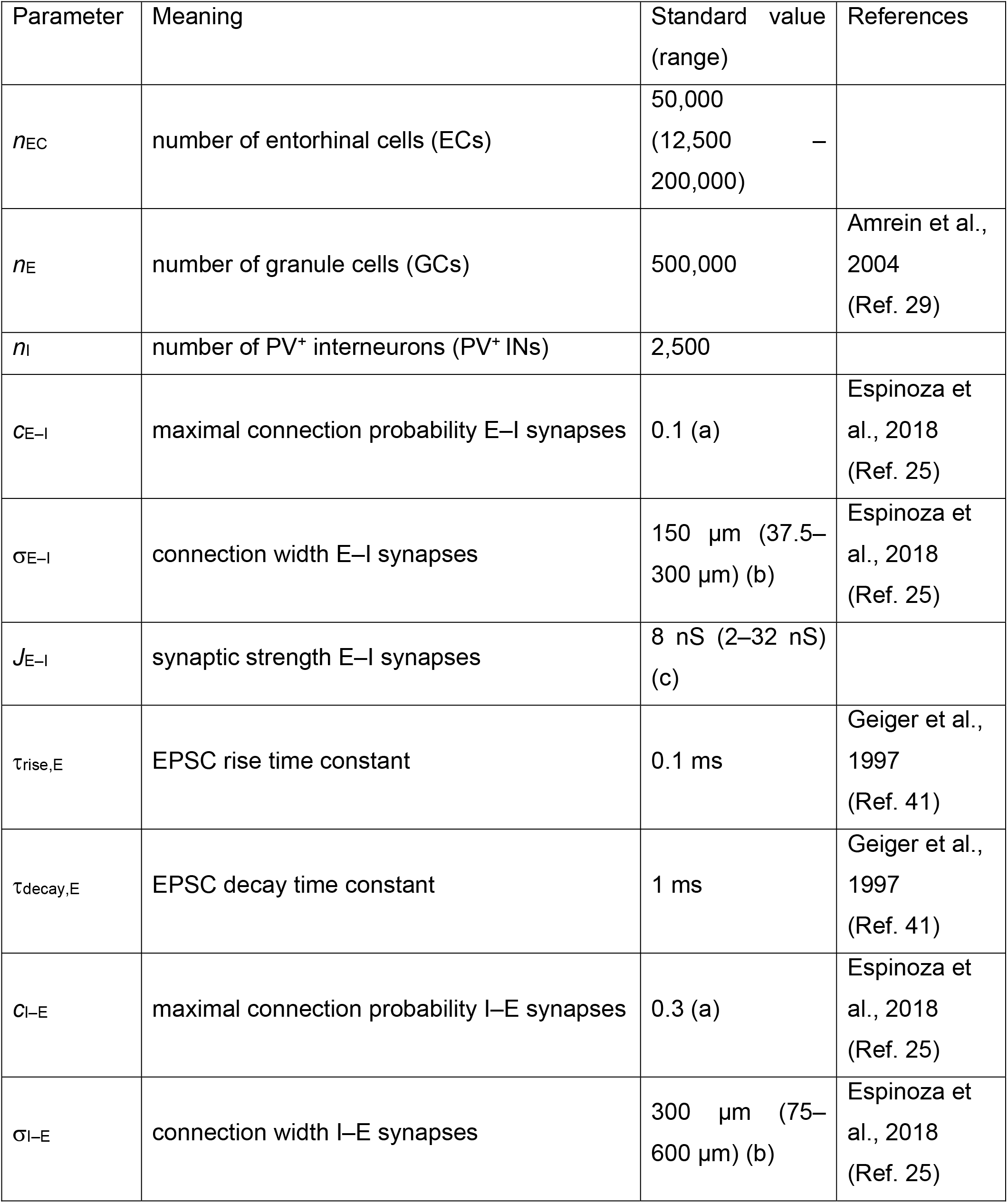
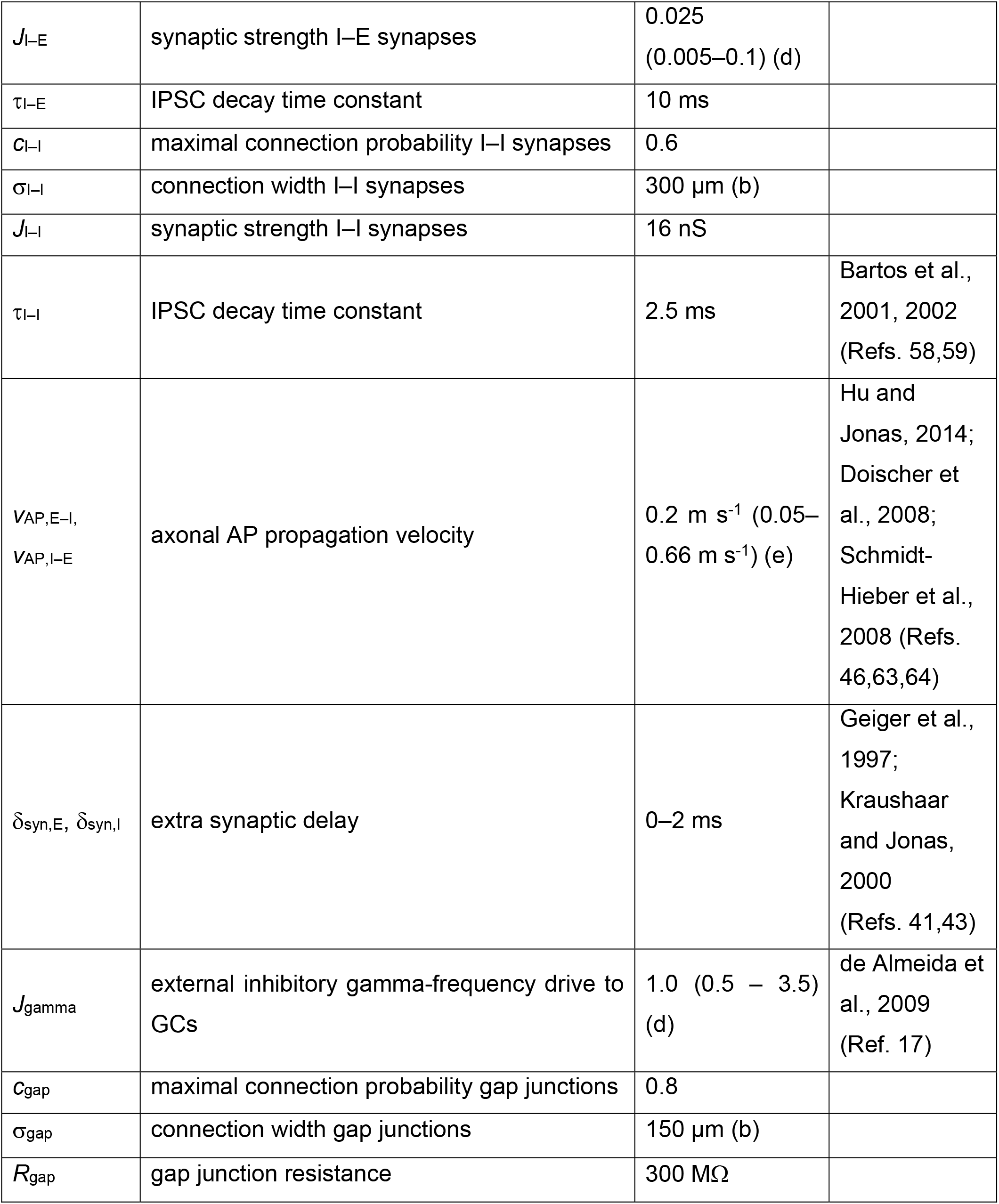
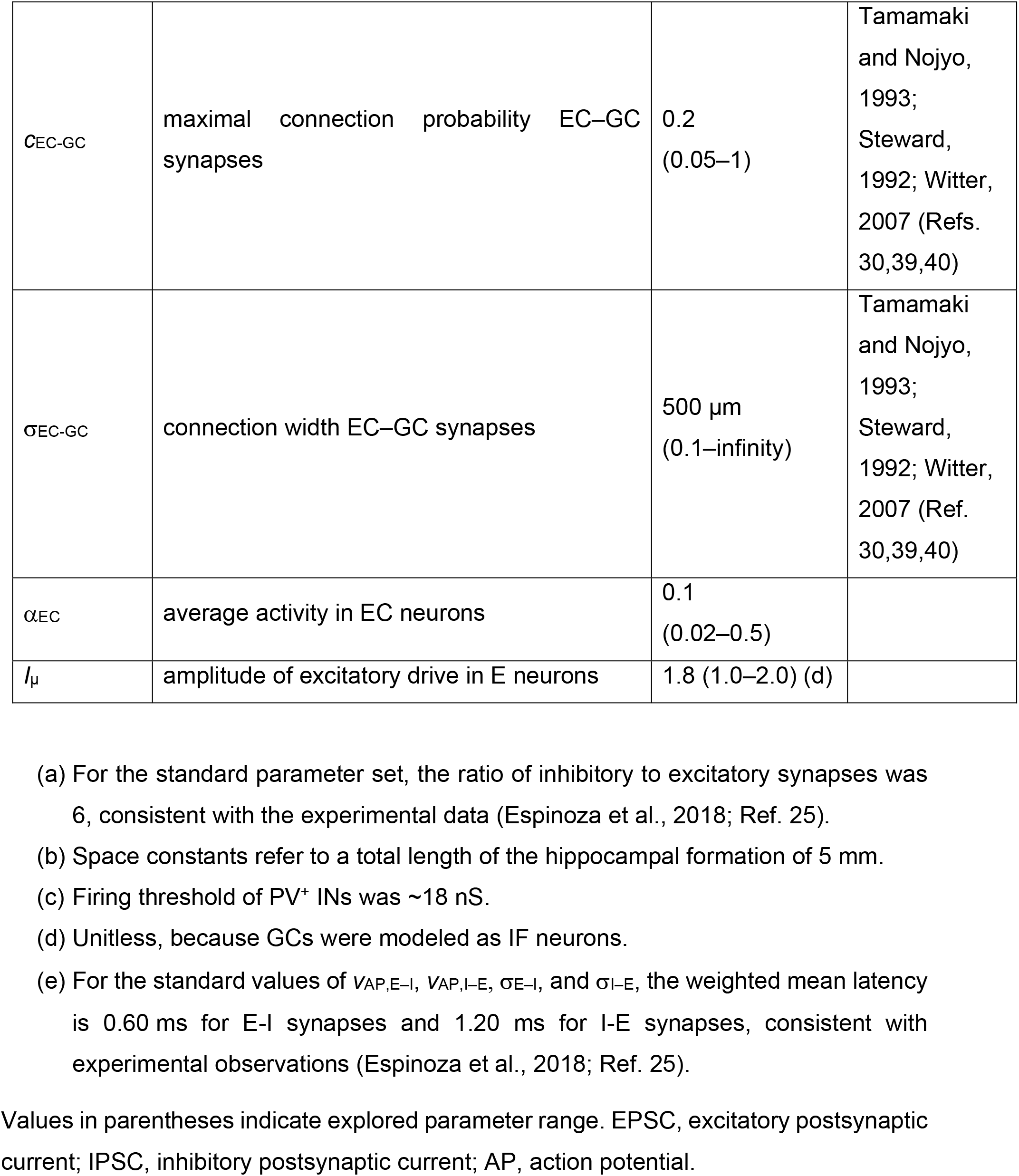
Standard parameters for the full-scale network model of pattern separation.

**Fig. 2.**
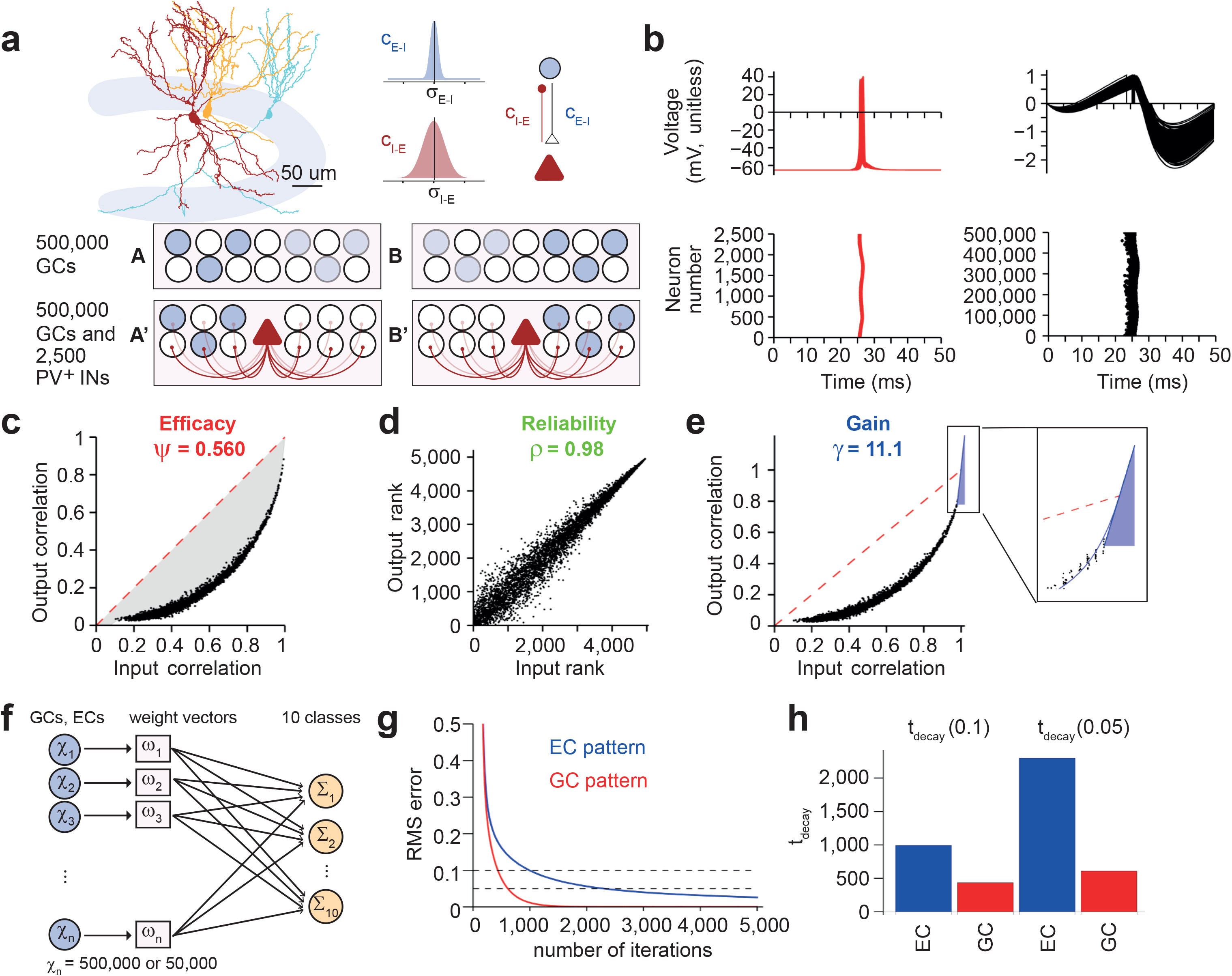
A biologically realistic PN–IN network including a winner-takes-all mechanism by lateral inhibition generates efficient pattern separation. (**a**) Top, illustration of experimentally determined connectivity rules between GCs and PV^+^ interneurons in dentate gyrus. Reconstruction was obtained from Espinoza et al., 2018 (Ref. 25). Bottom, schematic illustration of network mechanisms of pattern separation by a winner-takes-all mechanism. For simplicity, only GCs (circles) and INs (triangles) are depicted. Blue indicates cell activity, with color intensity reflecting the level of excitatory synaptic drive. Left, activity pattern A, right, activity pattern B. Top, network in the absence of inhibition, bottom, network in the presence of inhibition. (**b**) Top, membrane potential in INs (left, red) and PNs (right, black). Traces from every 10^th^ interneuron (250 traces total) and every 1,000^th^ GC (500 traces total) are superimposed. For GCs, membrane potential is unitless, since GCs were simulated as IF neurons. Bottom, rasterplots of AP generation in INs (left, red) and GCs (right, black). Each point indicates an AP. *t* = 0 corresponds to onset of inhibitory conductance representing a gamma oscillation cycle in the network^17^. (**c–e**) Input–output function in a network with standard parameter settings. Data points represent pairwise correlation coefficients between input patterns (excitatory synaptic drive, *R*_in_) and corresponding output patterns (action potential activity, *R*_out_). Dashed red line indicates identity. With standard parameter settings, Ψ (determined from the area between data points and identity line) was 0.560, demonstrating efficient pattern separation (c). Furthermore, the reliability of pattern separation ρ, computed as the correlation of ranked *R*_out_ versus ranked *R*_in_ data, was close to 1 (d). Finally, the gain γ of pattern separation, determined from the maximal slope of a polynomial function fit to the data for x → 1, was 11.1 (e), demonstrating that the network amplifies small differences in the synaptic input patterns into large differences in the AP output patterns. Blue curve indicates fit function, blue line represents corresponding tangent. For details, see Methods. (**f**) Schematic illustration of the perceptron decoding method, used to address the relevance of pattern separation for downstream networks^11^. A cellular population of size *n*_cells_ is connected to 10 perceptrons. One-hundred patterns (*n*_patterns_ = 100) are randomly grouped into ten classes (*n*_classes_ = 10). Thus, there is one perceptron for each class. The weights of the connections are iteratively adjusted, so that the activity of the 10 perceptrons matches the classification of the patterns. Weights *w* are adjusted according to the difference between true and predicted classes (Δ_classes_ = *I*_r_ × A_patterns_ × Δ*y*, where *l*_r_ is learning rate, A_patterns_ is the pattern matrix of size *n*_cells_ × *n*_patterns_, and Δ*y* is the sum of squared differences between classification and prediction). (**g**)Plot of RMS (root mean square) error against the number of iterations in the perceptron decoder. The perceptron was trained either with the EC input pattern (blue; before pattern separation) or the GC activity pattern (red; after pattern separation in the dentate gyrus). RMS was calculated as the mean of squared differences between true and predicted classes. (**h**)Summary bar graph of error decay time (*t*_decay_, when RMS error reached 0.1 or 0.05) in the perceptron decoder. Blue, EC input pattern (before pattern separation); red, GC activity pattern (after pattern separation in the dentate gyrus). Note that the perceptron decoder learns more rapidly from the GC output patterns than the EC input patterns.

We then analyzed pattern separation in the biologically realistic, full-size network model. One-hundred correlated binary activity patterns were applied in ECs, and activity was simulated in GCs and interneurons (Fig. 2b; Supplementary Figure 1; Table 1). Whereas all interneurons generated spikes, the activity in the GC population was only 0.012, indicating sparse coding (Fig. 2b). Input-output correlation curves were located below the identity line, indicating efficient pattern separation in the model (Fig. 2c–e). For the standard network parameters, Ψ was 0.560, indicating a high efficacy of pattern separation (Fig. 2c). ρ was 0.98, implying a high reliability of the pattern separation process (Fig. 2d). Finally, γ was 11.1, suggesting a high gain of pattern separation, i.e. the ability to convert small differences in input patterns into large differences in output patterns (Fig. 2e). Similar results were obtained when the tonic EC–GC drive was replaced by a random train of fast excitatory synaptic waveforms of comparable strength (Supplementary Figure 2). Likewise, efficient pattern separation was also observed in a network model that incorporated feedforward activation of interneurons (Supplementary Figure 3). Finally, efficient pattern separation was observed in a network model with synaptic amplitude fluctuations, i.e. trial-to-trial (“type 1”) variability and synapse-to-synapse (“type 2”) variability (Supplementary Figure 4). In conclusion, a biologically realistic PN–IN network is able to efficiently and reliably perform pattern separation computations.

Pattern separation in the dentate gyrus may facilitate the storage and recall of information in downstream CA3 networks^2,5–7^. For example, pattern separation may avoid that correlated representations are confused or erased by catastrophic interference^18^. To test these predictions, we attached our dentate gyrus network model to a single-layer perceptron decoder endowed with backpropagation learning, intended to represent the CA3 network (Fig. 2f)^11,37^. We trained the perceptron decoder to divide patterns into 10 randomly assigned classes, and assessed the learning rate by plotting the classification error against the number of iterations. To assess the effects of pattern separation, we compared the learning rates of the perceptron decoder for “unprocessed” EC patterns and “processed” GC patterns. Remarkably, the learning rate of the perceptron decoder was substantially faster for the GC patterns than for the corresponding EC patterns (Fig. 2g, h). These results demonstrate that the decorrelation generated by pattern separation in the dentate gyrus can be beneficial for computations in downstream networks, resulting in an improvement in the storage of information.

### Lateral inhibition is a primary mechanism underlying pattern separation

To identify the key mechanisms underlying pattern separation in the network model, we systematically varied the biologically relevant parameters (Fig. 3). First, we changed the amplitude of the excitatory synaptic drive (*I*_µ_) and the inhibitory gamma input (*J*_gamma_) in the network, parameters expected to affect thresholding properties of input-output conversion (Fig. 3a). Pattern separation was highly dependent on both parameters. Contour plot analysis revealed that the combination of small excitatory synaptic drive with small gamma input provided efficient pattern separation (Fig. 3b). As the excitatory drive was increased, a higher inhibitory gamma input was required to maintain the efficacy of pattern separation. Thus, the balance between excitatory drive and inhibitory gamma input determined the efficacy of pattern separation.

**Fig. 3.**
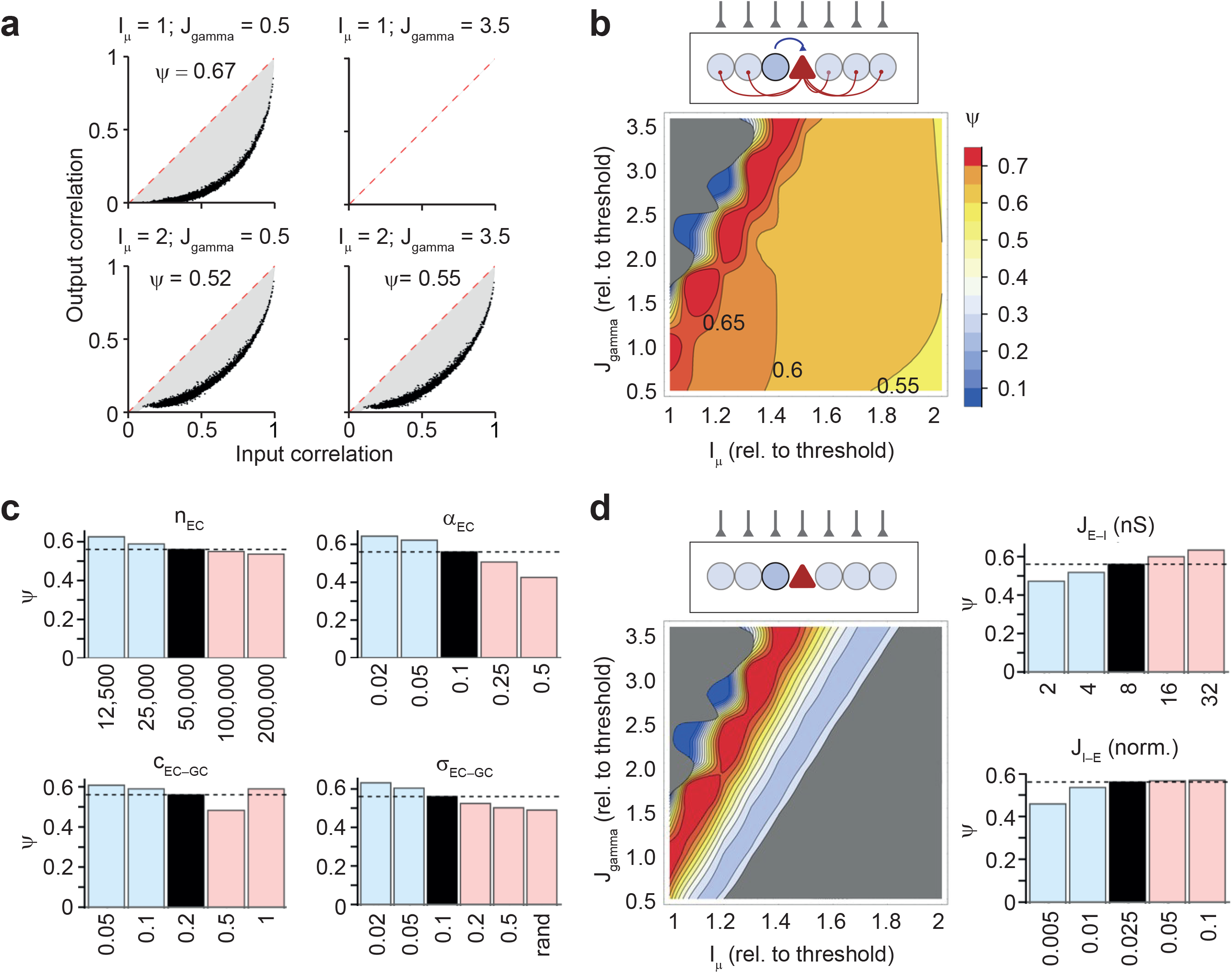
A winner-takes-all mechanism by lateral inhibition plays a critical role in pattern separation. (**a**) Input–output correlation plots for variations in network parameters in comparison to default values. Top, *I*_µ_ was set to 1, while external inhibitory gamma drive *J*_gamma_was set to 0.5 (left) or 3.5 (right). Bottom, similar to top, but *I*_µ_ was set to 2. Note that efficient pattern separation was observed in all scenarios except the condition with low excitatory drive and high *J*_gamma_(where activity α was 0). (**b**)Contour plot of Ψ against the mean excitatory drive (*I*_µ_, abscissa) and amplitude of external inhibitory gamma drive (*J*_gamma_, ordinate). Contour lines indicate Ψ; warm colors represent high values, whereas cold colors indicate low values. In the gray part of the plotting range, reliability was ρ < 0.1, or activity α was > 0.8 (i.e. majority of cells were firing). Note efficient pattern separation in a large subregion of the *I*_µ_–*J*_gamma_parameter space. (**c**)Divergent connectivity between ECs and GCs is important for efficient pattern separation. Effects of changes in number of ECs (*n*_EC_, top left), average activity of ECs (α_EC_, top right), maximal connection probability of EC–GC connectivity (*c*_EC–GC_, bottom left), and width of EC–GC connectivity (σ_EC–GC_, bottom right). Note that low EC number (*n*_EC_), sparse EC activity (α_EC_), and sparse EC–GC connectivity facilitate efficient pattern separation. (**d**)Lateral inhibition is necessary for efficient pattern separation. Left, contour plot of Ψ against the mean excitatory drive (*I*_µ_, abscissa) and amplitude of external inhibitory gamma drive (*J*_gamma_, ordinate) after complete elimination of lateral inhibition (*c*_E–I_= 0 and *c*_I–E_ = 0). Note efficient pattern separation in only a minimal subregion of the *I*_µ_–*J*_gamma_ parameter space. Right, effects of changes in synaptic strength of excitatory E–I synapses (*J*_E–I_) and inhibitory I–E synapses (*J*_E–I_). Black bars indicate Ψ for standard parameter settings; light blue bars represent reduced values; light red bars indicate increased values in comparison to standard values. Note that reduction in either *J*_E–I_or *J*_E–I_reduces the Ψ value.

Next, we determined how the properties of the synaptic input from ECs via the perforant path determined pattern separation^9–11,38^. To address this, we varied the number of entorhinal cells (*n*_EC_), the average EC activity level (α_EC_), and peak value and width of EC-GC connectivity (*c*_EC–GC_and σ_EC–GC_; Fig. 3c)^30,39,40^. Increasing the number of ECs decreased Ψ, whereas decreasing the number increased it (Fig. 3c, top left). Likewise, increasing the average EC activity decreased Ψ, whereas decreasing the activity had the reverse effect (Fig. 3c, top, right). Furthermore, increasing the EC–GC connection probability and the width mostly decreased Ψ, whereas decreasing probability or width led to opposite changes (Fig. 3c, bottom; Supplementary Figure 5). Effects of connection probability and width were similar when the GC drive values were randomly shuffled, indicating that spatial correlations in the input played only a minor role in pattern separation (Supplementary Figure 6). Interestingly, the effects of the *n*_EC_: *n*_GC_ ratio and *c*_EC–GC_ remained relatively minor even when the parameters were varied over a much wider range in a simplified system comprised of ECs, GCs, and a winner-takes-all mechanism in which the threshold was set according to the specified activity level (Supplementary Figure 7). Thus, the properties of the excitatory synaptic input influence pattern separation, but quantitatively play a relatively minor role.

Finally, we tested the contribution of lateral inhibition to pattern separation in the network model (Fig. 3d). Complete elimination of both excitatory E–I and inhibitory I–E synapses severely impaired pattern separation. Contour plot analysis of Ψ against *I*_µ_ and *J*_gamma_in the absence of lateral inhibition revealed Ψ values > 0.5 were only obtained in a small part of the parameter space (Fig. 3d, left). Furthermore, reducing the strength of either excitatory E–I or inhibitory I–E connections (*J*_E–I_ or *J*_I–E_) substantially reduced pattern separation efficacy Ψ (Fig. 3d, right). Similarly, reducing the peak connectivity or connectivity width of either excitatory E–I or inhibitory I–E connections (*c*_E–I_ and σ_E–I_, *c*_I–E_ and σ_I–E_) markedly affected pattern separation (Supplementary Figure 8). Thus, interfering with disynaptic inhibition at multiple levels uniformly decreased the efficacy of pattern separation. Taken together, these results indicate that lateral inhibition plays an essential role in pattern separation.

### Fast signaling and focal connectivity of PV^+^ interneurons are necessary for efficient pattern separation

If lateral inhibition plays a key role for pattern separation in the network, how do functional properties and connectivity rules affect this process? A hallmark property of PV^+^ GABAergic interneurons is their fast signaling at the level of synaptic input, input-output conversion, and synaptic output^26,41–43^. To test whether these fast signaling properties are relevant for pattern separation, we systematically varied the corresponding parameters in the model (Fig. 4a, b). Increasing the synaptic delay at excitatory GC–PV^+^ interneuron input synapses markedly impaired pattern separation (Fig. 4a, b, top left). Similarly, prolonging the time constants of the synaptic currents at excitatory GC–PV^+^ interneuron synapses reduced pattern separation performance (Fig. 4a, b, top right). Furthermore, increasing the membrane time constant of the PV^+^ interneurons reduced pattern separation performance (Fig. 4a, b, bottom left). Finally, increasing the synaptic delay at inhibitory PV^+^ interneuron–GC output synapses substantially impaired pattern separation (Fig. 4a, b, bottom right; Supplementary Figure 9). Thus, the fast signaling properties of PV^+^ interneurons are critical for the pattern separation process.

**Fig. 4.**
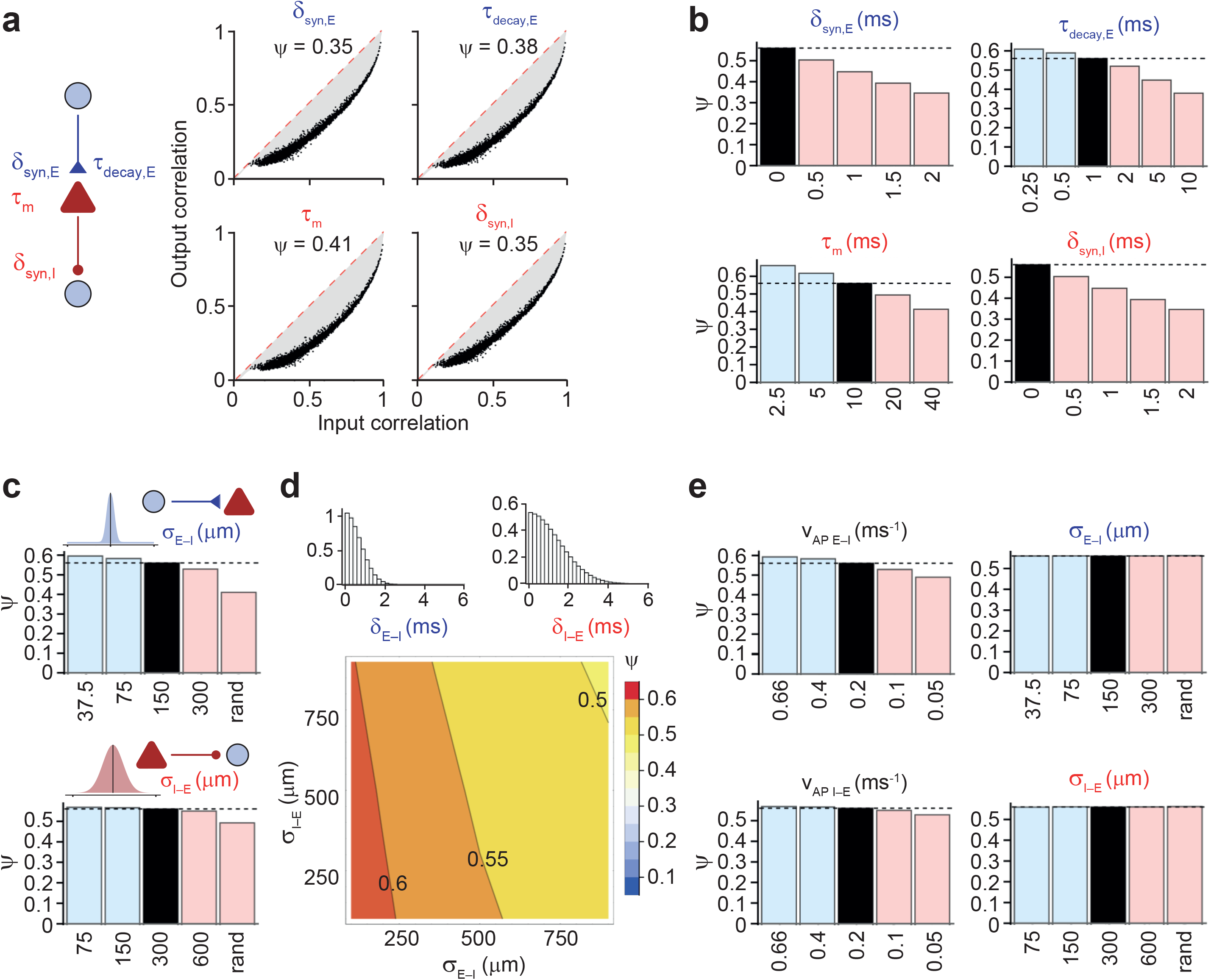
Fast interneuron signaling and focal PN–IN interconnectivity synergistically enhance pattern separation performance. (**a**) Effects of impairment of fast interneuron signaling on input-output correlation plots. Top left, introduction of an additional delay at GC–PV^+^ interneuron synapses (δ_syn,E_ = 2 ms); top right, prolongation of rise and decay time constant of synaptic conductance change at GC–PV^+^ interneuron synapses (rise time constant τ_rise,E_ = 1 ms; decay time constant τ_decay,E_ = 10 ms); bottom left, prolongation of the membrane time constant in PV^+^ interneurons (τ_m_= 40 ms); bottom, right, introduction of an additional delay at PV^+^ interneuron–GC synapses (δ_syn,I_ = 2 ms). (**b**) Summary bar graph of pattern separation efficiency Ψ for impairment of fast interneuron signaling for changes in δ_syn,E_ (top, left), τ_decay,E_ (top, right), τ_m_(bottom, left), and δ_syn,I_ (bottom, right). Interfering with fast signaling at multiple levels of the lateral inhibition pathway convergently impairs pattern separation efficacy. (**c**) Effects of focal connectivity on pattern separation. Summary bar graph of Ψ for different values of excitatory σ_E–I_ (top) or inhibitory σ_I–E_ (bottom) connectivity in the network. Right bar in each bar graph (“rand”) represents uniform random connectivity. Peak connectivity (and, if required, synaptic strength) was compensated to maintain the total synaptic efficacy. (**d**)Contour plot of Ψ against width of excitatory E–I connectivity (σ_E–I_) and inhibitory connectivity (σ_I–E_). Peak connectivity was compensated, as in (c). Note that networks with focal connectivity show more efficient pattern separation than networks with broad connectivity. Furthermore, asymmetry in spatial connectivity rules supports pattern separation. This is consistent with experimental observation of focal excitatory E–I connectivity versus the broader inhibitory I–E connectivity^25^. Inset on top, distribution of synaptic latency values in the network with standard parameters for excitatory E–I and inhibitory I–E synapses (probability density functions). (**e**) Effects of focal connectivity are mediated via effects on signaling speed. Left, summary bar graph of Ψ for different AP propagation velocity values for excitatory GC–PV^+^ interneuron synapses (*v*_AP,E–I_, top) and inhibitory PV^+^ interneuron–GC synapses (*v*_AP,I–E_, bottom). Right, summary bar graph of Ψ for different values of excitatory σ_E–I_(top) or inhibitory σ_I–E_ (bottom) connectivity after compensatory adjustment of both connectivity and delay to maintain both total connectivity and average delay at their default values. Note that broadening of connectivity fails to reduce pattern separation performance in the presence of delay adjustment. Thus, the beneficial effects of focal connectivity are largely generated by faster signaling.

The high pattern separation efficacy observed in the network model was surprising, because the model contains focal connectivity rules for both excitatory E–I and inhibitory I–E synapses in dentate gyrus^25^. In contrast, an efficient winner-takes-all mechanism may require lateral inhibition with long-range connectivity to ensure that a winner suppress all non-winners in the network. To resolve this apparent contradiction, we explored the effects of focal E–I and I–E connectivity in the network model (Fig. 4c–e). To address the effects of focal connectivity in isolation, we maintained the total connectivity (i.e. the area under the connection probability–distance curve) through compensatory changes of maximal connection probability (Fig. 4c). Increasing the width of connectivity for either excitatory E–I or inhibitory I–E synaptic connections reduced Ψ; particularly large changes were observed when focal connectivity was fully replaced by global random connectivity (Fig. 4c; Supplementary Figure 9). Thus, focal PN–IN connectivity supported pattern separation more effectively than global connectivity.

Next, we examined the effects of combined changes in the width of excitatory E– I and inhibitory I–E connectivity (Fig. 4d). As in the previous set of simulations, we maintained the total connectivity. Contour plot analysis confirmed that focal connectivity supported pattern separation more effectively than broad connectivity. However, the effects of changes in the width of excitatory E–I and inhibitory I–E connectivity were asymmetric. Thus, a high Ψ was obtained in a configuration in which the excitatory E–I was more focal than the inhibitory I–E connectivity (Fig. 4d). This was consistent with experimental observations that excitatory E–I is more focal than inhibitory I–E connectivity and that lateral inhibition is highly abundant in the circuit^25^. In conclusion, lateral inhibition effectively supported pattern separation.

Why does focal connectivity support pattern separation better than global connectivity? One possibility is that the effects of focal connectivity might be a consequence of changes in average latency, which are shorter in a focally connected network than in an equivalent random network. To test this hypothesis, we examined the effects of changes in axonal AP propagation velocity at excitatory E–I and inhibitory I–E synapses on pattern separation. Slowing AP propagation reduced Ψ, whereas accelerating propagation increased it (Fig. 4e, left). To test whether changes in synaptic latency fully account for the functional differences between focal and random networks, we changed the connectivity width while maintaining the kinetic properties of disynaptic inhibition through compensatory changes of AP propagation velocity (Fig. 4e, right). Notably, changes in propagation velocity almost completely compensated the effects of changes in connectivity. Thus, focal connectivity and fast biophysical signaling in GC– PV^+^ interneuron microcircuits play synergistic roles in providing rapid lateral inhibition, an essential requirement for efficient pattern separation.

## DISCUSSION

A fundamental question in neuroscience is how the properties of synapses and microcircuits contribute to higher-order computations in the brain. Our network model provides some answers to this central question, for a specific network function (pattern separation) and a specific circuit (dentate gyrus). First, our results provide a proof-of-principle that a biologically realistic network model is a highly efficient pattern separator. Second, our results show that lateral inhibition plays a critical role in the pattern separation process. Finally, they indicate that fast biophysical signaling properties of PV^+^ interneurons and focal connectivity are essential for efficient pattern separation.

Previous work in the cerebellum suggested that expansion of coding space is a key mechanism underlying pattern separation^9–11^. Our computational analysis confirms that the connectivity rules between ECs and GCs play an important role in pattern separation. First, the number of ECs is relevant, with a smaller number of neurons resulting in more efficient pattern separation (Fig. 3c). This is consistent with previous models, which emphasized the role of code expansion^9–11,38^. Second, the average EC– GC connectivity is important, with sparse connectivity enhancing pattern separation performance (Fig. 3c). Although this is also true for the cerebellum^11^, the mechanisms may be different in the hippocampus, because GCs receive a much higher number of synaptic inputs (> 1,000)^31,32^ compared to GCs in cerebellum (~5)^11^. Finally, a mix of structured and random EC–GC connectivity is optimal for the pattern separation mechanism (Supplementary Figure 6). However, the effects of these parameters on pattern separation efficacy are moderate. Thus, the rules of EC–GC connectivity, although clearly important, are not the main determinants of pattern separation in the dentate gyrus.

Previous studies suggested a major role of inhibition in pattern separation in the olfactory bulb of mammals and zebrafish and in the equivalent mushroom body of Drosophila^18–20^. Furthermore, a role of inhibition has been suggested in the hippocampus^24,44^. Recent functional connectivity analysis between GCs and interneurons revealed that lateral inhibition is uniquely abundant in the dentate gyrus^25^. Here, we show that lateral inhibition inserted into a biologically inspired network model generates a powerful winner-takes-all mechanism. Both excitatory E–I synapses and inhibitory I–E synapses are necessary for pattern separation (Fig. 3d). Remarkably, the winner-takes-all mechanism based on lateral inhibition works in a network comprised of a relatively small number of neurons. Winner-takes-all computations are also performed by networks of perceptrons^16^. However, in such implementations, pattern separation requires a multi-layer structure with a much larger number of neuron-like elements and synaptic connections^16^. Thus, lateral inhibition represents a compact, resource-efficient implementation of a winner-takes-all computation.

Our results reveal two novel determinants of the efficacy of pattern separation. The first key factor is fast signaling in GABAergic cells. This may have been expected, because sufficient speed is required to ensure that a small number of winners suppresses a large number of non-winners (Fig. 4a, b). Lateral inhibition in the dentate gyrus is primarily mediated by PV^+^ interneurons, since these interneurons are more connected than other interneurons (such as somatostatin^+^ or cholecystokinin^+^ interneurons)^25,45^. Furthermore, PV^+^ interneurons express an extensive repertoire of fast biophysical signaling mechanisms at the level of synaptic input, AP initiation, and synaptic output^26,41,46^. Thus, PV^+^ interneurons are prime candidates for the neuronal implementation of a winner-takes-all mechanism by lateral inhibition. However, the contribution of other interneuron subtypes cannot be excluded.

The second key factor is focal connectivity between principal neurons and interneurons, which substantially enhances pattern separation. This is counter-intuitive, because a long-range divergent output may be useful to suppress all non-winners ^15,16^. However, our simulations show that networks with focal connectivity are more effective than networks with wide connectivity (Fig. 4c). Furthermore, the pattern separation mechanism works well if the connectivity is asymmetric, with excitatory E–I synapses showing narrower connectivity and inhibitory I–E synapses wider connectivity, as observed experimentally (Fig. 4d)^25^.

Our results demonstrate that pattern separation can accelerate learning by a downstream perceptron decoder (Fig. 2f–h). Does this also happen in the biological network? Hippocampal GCs connect to CA3 pyramidal neurons via hippocampal mossy fiber synapses^5^. Because of their large size, these synapses are often viewed as “detonator” synapses^47^. If the mossy fiber synapses are detonators, one would expect that the decorrelated signals are relayed to the CA3 network, and trigger efficient storage of information in CA3–CA3 synapses, similar to the perceptron^6,7^. However, recent work suggests that the signaling properties of mossy fiber synapses are more complex, since subdetonation and conditional detonation can coexist with plasticity-dependent full detonation^48^. If the mossy fiber output would be slightly below the detonation threshold, this may introduce another mechanism of synaptic integration and thresholding into the network, which could amplify the degree of pattern separation. Large-scale network simulations including both dentate gyrus and CA3 will be needed to further address this possibility.

Taken together, the present results add to the emerging view that PV^+^ interneurons are not only involved in basic microcircuit functions, such as feedforward and feedback inhibition, but also contribute to higher-order computations in neuronal networks^26^. Consistent with this idea, pharmacological analysis revealed that inhibition plays a role in pattern separation in behavioral experiments^44^. More specific optogenetic and pharmacogenetic strategies will be needed to further delineate the contribution of PV^+^ interneurons and other interneurons to these processes. Finally, since accumulating evidence suggests that PV^+^ interneuron dysfunction is associated with brain disorders, including schizophrenia^26^, it will be important to evaluate whether pattern separation is impaired and how exactly inhibition contributes to circuit dysfunction in these diseases^49^.

## METHODS

### Topology of a full-size dentate gyrus network model

The pattern separation network model consists of two layers, the first layer representing the entorhinal cortex, with 50,000 ECs, and the second layer representing the dentate gyrus, with 500,000 GCs and 2,500 PV^+^ interneurons (INs). First and second layer were connected by EC–GC synapses, representing the perforant path input to the dentate gyrus. A winner-takes-all mechanism mediated by lateral inhibition was implemented by connecting GCs and INs by excitatory E–I synapses in one direction and by inhibitory I– E synapses in the other direction.

Unlike other models of dentate gyrus circuits^24,50^, the model was implemented in full size. The number of GCs was chosen to represent the dentate gyrus of one hemisphere in adult laboratory mice^29^. Full-scale implementation was necessary: (1) to increase the realism of the simulations, (2) to be able to implement measured macroscopic connectivity rules without scaling^51^, and (3) to simulate sparse coding regimes, which were unstable in smaller networks (Fig. 1e).

The model was designed to incorporate the connectivity rules of PV^+^ interneurons and GCs in the dentate gyrus^25^. Other types of interneurons, such as SST^+^ hilar interneurons with axons associated with the perforant path or CCK^+^ hilar interneurons with axons associated with the commissural / associational pathway^45,52–54^, were not explicitly included because of their low connectivity^25^ and their slower signaling speed^26^. While the first property of SST^+^ or CCK^+^ interneurons would make them less likely to be activated by GC activity, the second property would make them less suitable for the neuronal implementation of a winner-takes-all mechanism^17^. In total, the conclusions of the present paper were based on 594 full-scale simulations.

### Implementation of inhibitory interneurons

Interneurons were implemented as single-compartment, conductance-based neurons to capture the electrical properties of PV^+^ interneurons. Membrane potential was simulated by solving the equation:

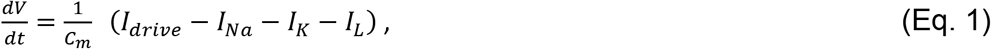

where *V* is membrane potential, *t* is time, *C*_m_ is membrane capacitance, *I*_drive_ is driving current, while *I*_Na_, *I*_K_, and *I*_*L*_represent sodium, potassium and leakage current, respectively. *I*_Na_was modeled as

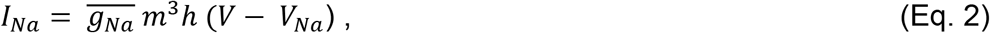

where 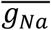 is the maximal sodium conductance, *m* is the activation parameter, *h* is the inactivation parameter, and *V*_Na_represents the sodium ion equilibrium potential.

Similarly, *I*_K_ was modeled according to the equation

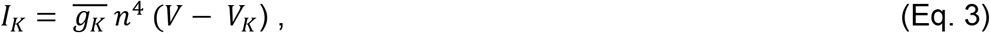

where 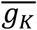 is the maximal potassium conductance, *n* is the activation parameter, and *V*_K_ represents the potassium ion equilibrium potential.

Finally, *I*_*L*_ was given as

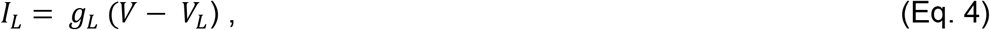

where *g*_L_ is leakage conductance and *V*_L_ is corresponding reversal potential.

State parameters *m*, *h*, and *n* were computed according to the differential equation

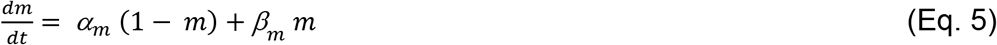

and equivalent equations for *h* and *n*.

α_m_, α_h_, α_n_ values and β_m_, β_h_, β_n_ values were calculated according to the equations α_m_= 0.1 ms^-1^ × −(*V*+35 mV) / {Exp[−(*V*+35 mV)/10 mV] – 1}, β_m_ = 4 ms^-1^ × Exp[−(*V*+60 mV)/18 mV], α_h_ = 0.35 ms^-1^ × Exp[−(*V*+58 mV)/20 mV], β_h_= 5 ms^-1^ / {Exp[−(*V*+28 mV)/10 mV] + 1}, α_n_ = 0.05 ms^-1^ × −(*V*+34 mV) / {Exp[−(*V*+34 mV)/10 mV] − 1}, and β_n_ = 0.625 ms^-1^ × Exp[−(*V*+44 mV)/80 mV]^55^. Single neurons were assumed to be cylinders with diameter and length of 70 µm, giving a surface area of 15,394 µm^2^ and an input resistance of 65 MΩ^42^. Neurons showed a rheobase of 39 pA and a fast-spiking, type I AP phenotype^56^, as characteristic for PV^+^ interneurons^26^. Maximal conductance values 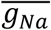, 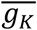, and *g*_L_ were set to 35 mS cm^−2^, 9 mS cm^−2^, and 0.1 mS cm^−2^, respectively^55^. *V*_Na_ and *V*_K_ equilibrium potentials were assumed as 55 mV and −90 mV, respectively. Finally, *V*_L_ was set to −65 mV.

### Implementation of GCs

GCs were implemented as leaky integrate-and-fire (IF) spiking neurons. To enable the integration of excitatory and inhibitory synaptic events with different kinetics, the standard IF model was extended as follows^57^:

The time course of synaptic excitation was described by the differential equation

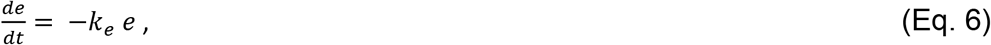

where *k*_e_ is the synaptic excitation rate constant, i.e. the inverse of the time constant.

Likewise, the time course of synaptic inhibition was described by the differential equation

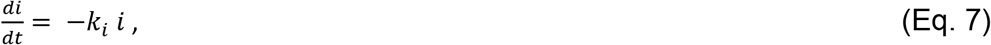

where *k*_i_ is the synaptic inhibition rate constant.

Finally, the firing of the neuron was controlled by a membrane state variable *v*; when *v* reaches one, the cell fires, which resets the membrane by returning *v* to 0. The time course of *v* was determined by the differential equation

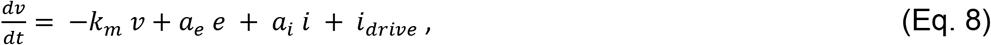

where *k*_m_is inverse of the membrane time constant, a_e_ and a_i_ are amplitudes of synaptic events, and *i*_drive_ represents the excitatory drive any given neuron receives^57^. Excitation time constant, inhibition time constant, and membrane time constant were set to 3, 10, and 15 ms, respectively^25,32,43^. The refractory period was assumed as 5 ms. Note that in the IF model *v*, *e*, *i*, and *i*_drive_ are unitless.

### Implementation of synaptic interconnectivity

Synapses between neurons were placed with distance-dependent probability. Normalized distance was cyclically measured as

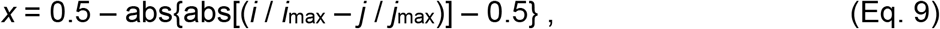

where *i* and *j* are indices of pre- and postsynaptic neurons, *i*_max_ and *j*_max_ are corresponding maximum index values, and abs(r) is the absolute value of a real number r. Connection probability was then computed with a Gaussian function as

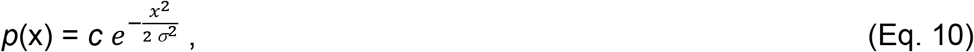

where *c* is maximal connection probability (*c*_E–I_, *c*_I–E_, *c*_I–I_, and *c*_gap_, respectively) and σ is the standard deviation representing the width of the distribution (σ_E–I_, σ_I–E_, σ_I–I_, and σ_gap_; Table 1).

Connection probability between ECs and GCs was computed from a Gaussian function with peak connection probability of 0.2 and a standard deviation of 500 µm, to represent the divergent connectivity from the entorhinal cortex to the dentate gyrus^30,39,40^. Binary activity patterns in upstream ECs were converted into patterns of excitatory drive of GCs. Although this drive was primarily intended to represent input from entorhinal cortex neurons, it may include contributions from other types of excitatory neurons (e.g. mossy cells or CA3 pyramidal cells)^50^.

Excitatory GC–interneuron synapses, inhibitory interneuron–GC synapses, and inhibitory interneuron–interneuron synapses were incorporated by random placement of NetCon objects in NEURON^57^; gap junctions were implemented by random placement of pairs of point processes. For excitatory GC–interneuron synapses and inhibitory interneuron–interneuron synapses, synaptic events were simulated using the Exp2Syn class of NEURON. For excitatory GC–interneuron synapses, we assumed τ_rise,E_ = 0.1 ms, τ_decay,E_ = 1 ms, and a peak conductance of 8 nS^25,41^. For inhibitory interneuron– interneuron synapses, we chose τ_rise,I_ = 0.1 ms, τ_decay,I_ = 2.5 ms, and a peak conductance of 16 nS^25,58,59^. For inhibitory interneuron–GC synapses, the synaptic weight was chosen as 0.025 (unitless, because GCs were modelled as IF neurons). For all chemical synapses, synaptic latency was between 0 and 25 ms according to distance between pre- and postsynaptic neuron. Gap junction resistance was assumed as 300 MΩ, approximately five times the input resistance of the cell^25,58,59^. Synaptic reversal potentials were 0 mV for excitation and −65 mV for inhibition. The maximal length of the hippocampal network was assumed as 5 mm, consistent with anatomical descriptions in mice^60^.

### Detailed implementation and simulations

Simulations of network activity were performed using NEURON version 7.6.2^57^ in combination with Mathematica version 11.3.0.0 (Wolfram Research). Simulations were tested on reduced-size networks running on a PC using Windows 10. Full-size simulations were run on x86_64-based shared memory systems (Supermicro or SGI UV 3000 systems) using GNU/Linux (Debian, SLES).

Simulations were performed in four steps (Supplementary Figure 1). First, we computed random binary activity patterns in ECs. To generate input patterns with defined correlations over a wide range, 100 uncorrelated random vectors a_i_ of size *n*_EC_ were computed, where individual elements are pseudorandom real numbers in range of 0 to 1 and *n*_EC_ is the number of ECs. Vectors were transformed into correlated vectors as *r* × *a*_1_ + (1 − *r*) × *a*_i_, where *a*_1_ is the first random vector and *r* corresponds to the correlation coefficient. *r* was varied between 0.1 and 1. Finally, a threshold function *f*(x) = H(x − θ) was applied to the vectors, where H is the Heaviside function and θ is the threshold that determines the activity level in the pattern. Empirically, 100 input patterns were sufficient to continuously cover the chosen range of input correlations. Unless stated differently, the average activity in EC neurons (α_EC_), i.e. the proportion of spiking cells, was assumed to be 0.1.

Second, the patterns in the upstream neurons were converted into patterns of excitatory drive in GCs, by multiplying the activity vectors with the previously computed connectivity matrix between ECs and GCs. Unless otherwise indicated, the mean tonic current value was set to 1.8 times the threshold value of the GCs (i.e. *I*_µ_ = 1.8; unitless, since GCs were implemented as IF units; Table 1). In a subset of simulations (Supplementary Figure 2), the tonic current was replaced by Poisson trains of excitatory postsynaptic currents (EPSCs) to convey a higher degree of realism. In these simulations, events were simulated by NetStim processes. In another subset of simulations (Supplementary Figure 3), the tonic excitatory drive computed from the EC activity and the EC–GC connectivity was applied in parallel to GCs and INs after appropriate scaling to represent feedforward inhibition.

Third, we computed the activity of the network for all 100 patterns. Simulations were run with 5 µs fixed time step over a total duration of 50 ms. At the beginning of each simulation, random number generators were initialized with defined seeds to ensure reproducibility. At the beginning of each simulation, an inhibitory synaptic event of weight 1 (relative to threshold) was simulated in all GCs to mimic recovery from a preceding gamma cycle^17^. Spikes were detected when membrane potential reached a value of 1 in the GCs and 0 mV in the interneurons. Subsequently, spike times were displayed in raster plot representations. Furthermore, 100 binary output vectors were computed, by setting the value to 1 if a cell generated ≥ 1 spikes in the time interval 0 ≤ t ≤ 50 ms, and to 0 otherwise.

Finally, Pearson’s correlation coefficients were computed for all pairs of patterns 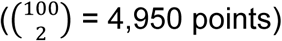, at both input (tonic excitatory drive vector) and output level (spike vector) in parallel, and output correlation coefficients (*R*_out_) were plotted against input correlation coefficients (*R*_in_). Pattern separation was quantitatively characterized by three parameters: (1) The efficacy of pattern separation (Ψ) was quantified by an integral-based index, defined as the area between the identity line and the *R*_out_ versus *R*_in_ curve, normalized by the area under the identity line 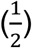. Thus,

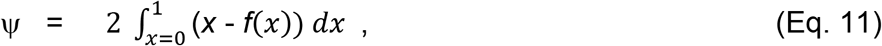

where *f*(x) represents the input-output correlation function. In practice, *f*(x) was determined by linear interpolation of data points after sorting by *R*_*in*_ values, averaging of points with same *R*_*in*_, and including points (0|0) and (1|1). Based on these definitions, a Ψ value close to 1 would correspond to an ideal pattern separator. In contrast, Ψ = 0 would represent pattern identity, whereas Ψ < 0 would indicate pattern completion^7^. (2) The reliability of pattern separation (ρ) was quantified by the Pearson’s correlation coefficient of the ranks of all *R*_*out*_ versus the ranks of all *R*_*in*_ data points. An ideal pattern separator will maintain the order of pairwise correlations: If a pair of patterns is more similar than another pair at the input level, it will be also more similar at the output level. Thus, for an ideal pattern separator, ρ will be close to 1. (3) Finally, the gain of pattern separation (γ) was quantified from the maximal slope of the *R*_out_ versus *R*_in_ curve. In practice, this value was determined from the first derivative of a 5^th^ or 10^th^-order polynomial function *f*(x) fit to the *R*_*out*_ versus *R*_*in*_ data points as 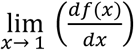; *f*(x) was constrained to pass through points (0|0) and (1|1). A γ value >> 1 would correspond to an ideal pattern separator. In contrast, γ = 1 would represent pattern identity, whereas γ < 1 would indicate pattern completion^7^.

### Analytical analysis of pattern separation

To describe the pattern separation process in a simple mathematical form (Fig. 1c, d), we obtained an analytical solution for the correlation coefficient of a bivariate Gaussian after dichotomization using Hoeffding’s lemma

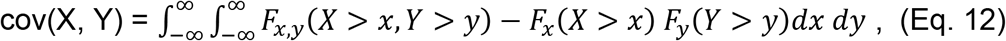

where cov is the covariance, *X* and *Y* are random variables, *F*_*x,y*_ denotes the joint probability function, and *F*_*x*_, *F*_*y*_ represent the marginal probability functions^28,61,62^. To simulate finite-size effects (Fig. 1e, f), vectors of real random numbers were drawn from a bivariate Gaussian distribution with defined correlation *R*_in_, converted into vectors of binary numbers by applying a threshold, and subjected to correlation analysis, resulting in the correlation coefficient *R*_out_. The threshold was chosen to reach a previously specified average activity level α, and the size of the vector varied in the range 5,000 to 50,000. Furthermore, in a subset of simulations (Supplementary Figure 7), activity was simulated in ECs, computed into drive patterns in GCs by multiplication with the EC–GC connectivity matrix, and directly converted into binary activity values in GCs by applying a threshold corresponding to the desired activity level α. This simplified approach permitted systematic variation of model parameters (e.g. cell numbers and connection probabilities) over a wide range.

### Analysis of input and output patterns by a perception decoder

To test whether the pattern separation process resulted in a gain of function that could be exploited by downstream networks, we analyzed input and output patterns by a perception decoder (Fig. 2f–h)^11,37^. The perception decoder was trained to categorize 100 input and output patterns into 10 random classes. The decoder was comprised of a single layer, and a backpropagation learning algorithm was used to iteratively adjust the weights. Initially, all weights were arbitrarily set to 0.1. The learning rate was assumed as 5 × 10^-4^. In each learning iteration, weights were adjusted according to the deviations between predicted and observed classifications. In total, 5,000 learning iterations were run, and the learning speed was quantified as the number of iterations at which the root mean square error reached a value of 0.1 or 0.05.

### Conventions

Throughout the paper, model parameters given in Table 1 are referred to as standard parameters. In summary bar graphs, black bars indicate these standard values, light blue bars reduced values, and light red bars increased values in comparison to the default parameter set. Throughout the paper, the term “pattern” is defined as a vector of real numbers (for excitatory drive) or a vector of binary values (for activity, 1 if the cell fires, 0 otherwise). In both cases, the vector length corresponds to the number of cells.

### Data and code availability

Original data, analysis programs, and computer code for network simulations will be provided by the corresponding author (P.J.) upon request. Simulation code will be updated according to new experimental information about connectivity (e.g. EC–GC connectivity rules). Furthermore, IF models of GCs and single-compartment models of interneurons will be gradually replaced by more detailed models (conductance-based models and multi-compartmental models, respectively).

## ACKNOWLEDGMENTS

We thank Drs. Ad Aertsen, Arnd Roth, and Federico Stella for critically reading earlier versions of the manuscript. We are grateful to Florian Marr and Christina Altmutter for excellent technical assistance, Eleftheria Kralli-Beller for manuscript editing, and the Scientific Service Units of IST Austria for efficient support. Finally, we thank Drs. Ted Carnevale, Laszlo Erdös, Michael Hines, Nancy Kopell, Duane Nykamp, and Dominik Schröder for useful discussions, and Rainer Friedrich and Simon Wiechert for sharing unpublished data. Parts of the results presented were obtained using the Mach2 Interuniversity Shared Memory Supercomputer (Linz, Austria). This project received funding from the European Research Council (ERC) under the European Union’s Horizon 2020 research and innovation programme (grant agreement No 692692) and the Fond zur Förderung der Wissenschaftlichen Forschung (Z 312-B27, Wittgenstein award), both to P.J.

## Author contributions

S.J.G. and P.J. designed the model and the layout of the simulations, A.S. performed large-scale simulations on computer clusters, C.E., X.Z., and B.A.S. provided experimental data, S.J.G. and P.J. analyzed data, and P.J. wrote the paper. All authors jointly revised the paper.

---

**Supplementary Figure 1.**
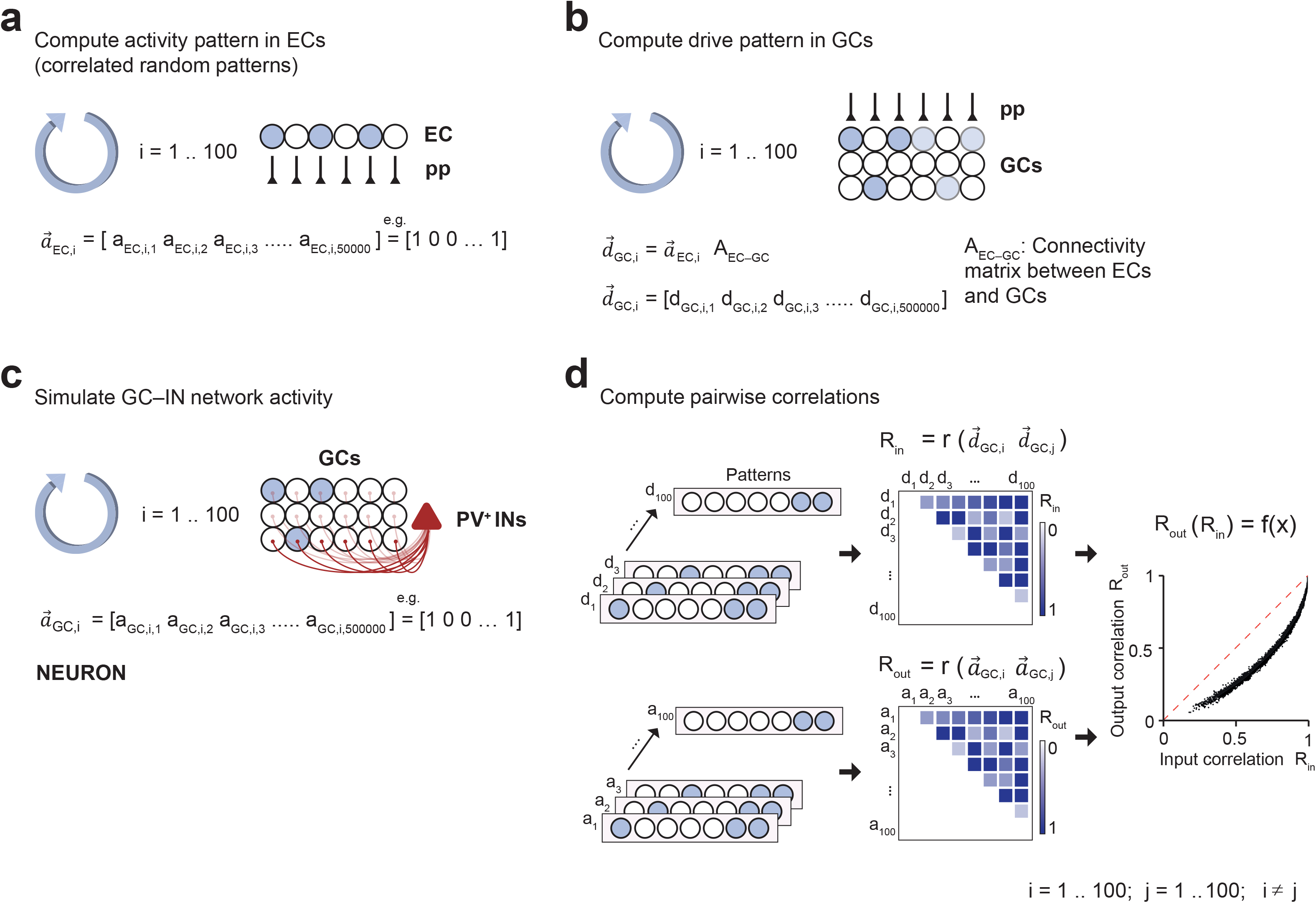
Schematic illustration of full-size network simulations. (**a**) Computation of activity in ECs. 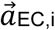 represents the i^th^ binary activity vector in the ECs (50,000 neurons). (**b**) Computation of drive patterns in GCs. 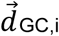 represents the i^th^ drive vector in GCs (500,000 neurons). 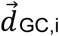 was computed as the product of activity vector 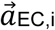 and connectivity matrix A_EC–GC_. (**c**)Computation of activity in dentate gyrus. Activity in the full-size network was simulated using NEURON version 7.6.2^57^. 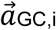 represents the i^th^ binary activity vector in the GCs, determined by the spiking of GCs. (**d**) Computation of pattern correlation and input-output correlation curves. Correlations *R*_*in*_ were computed between pairs of drive vectors, correlations *R*_*out*_ were computed between pairs of activity vectors. Finally, *R*_*out*_ and *R*_*in*_ values were plotted against each other, and a continuous function *f*(x) was obtained by linear interpolation.

**Supplementary Figure 2.**
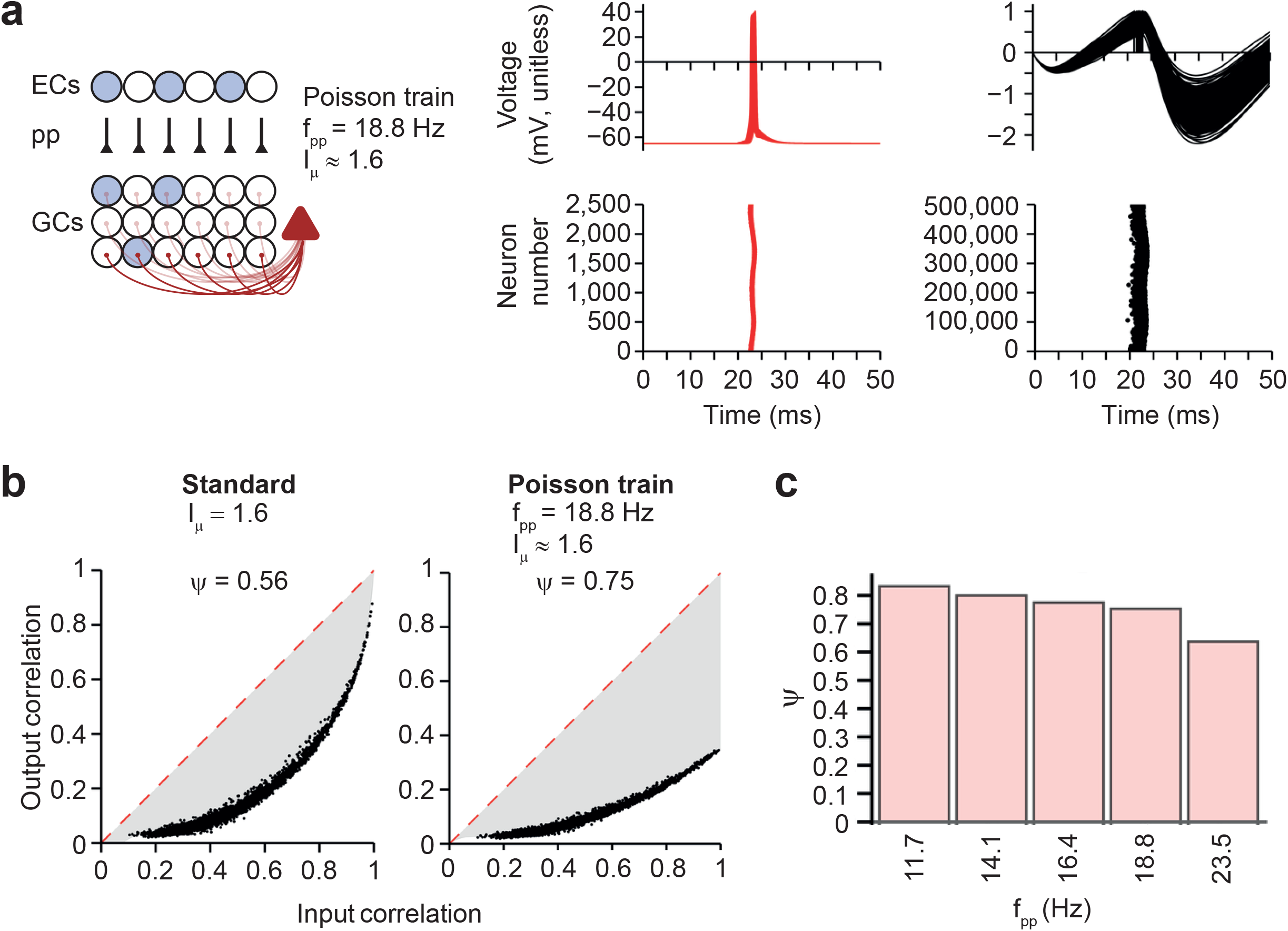
A network model activated by Poisson trains in perforant path input is able to perform efficient pattern separation. (**a**) Left, schematic illustration of network containing perforant path input. Right, simulated membrane potentials (top) and rasterplot of interneuron and principal neuron firing (bottom) in a model with realistic EC–GC synaptic input, represented by Poisson trains of APs at different frequency. Every point in the rasterplots represents an AP. Average activity frequency of the perforant path (pp) synapses *f*_pp_ = 18.8 Hz; activation frequency was chosen to give *I*_μ_ ≈ 1.6. (**b**) Input-output correlation curves for standard model with tonic excitatory drive (left; *I*_μ_ = 1.6) and model in which excitatory drive was generated by Poisson trains of EPSCs in GCs (right; *f*_pp_ = 18.8 Hz; activation frequency was chosen to give *I*_μ_ ≈ 1.6, facilitating the comparison with the standard model). Note that the randomness of the input trains resulted in a drop of the output correlation for input correlation values of 1, because an additional random process is added to the system. (**c**) Dependence of Ψ on activity frequency of perforant path synapses. Synaptic weight of EC–GC synapses was set to *J*_EC–GC_ = 0.002 in all simulations. Activation frequency was chosen to approximately match *I*_µ_ = 1, 1.2, 1.4, 1.6, and 2.0 in the standard model (see Fig. 3b and d).

**Supplementary Figure 3.**
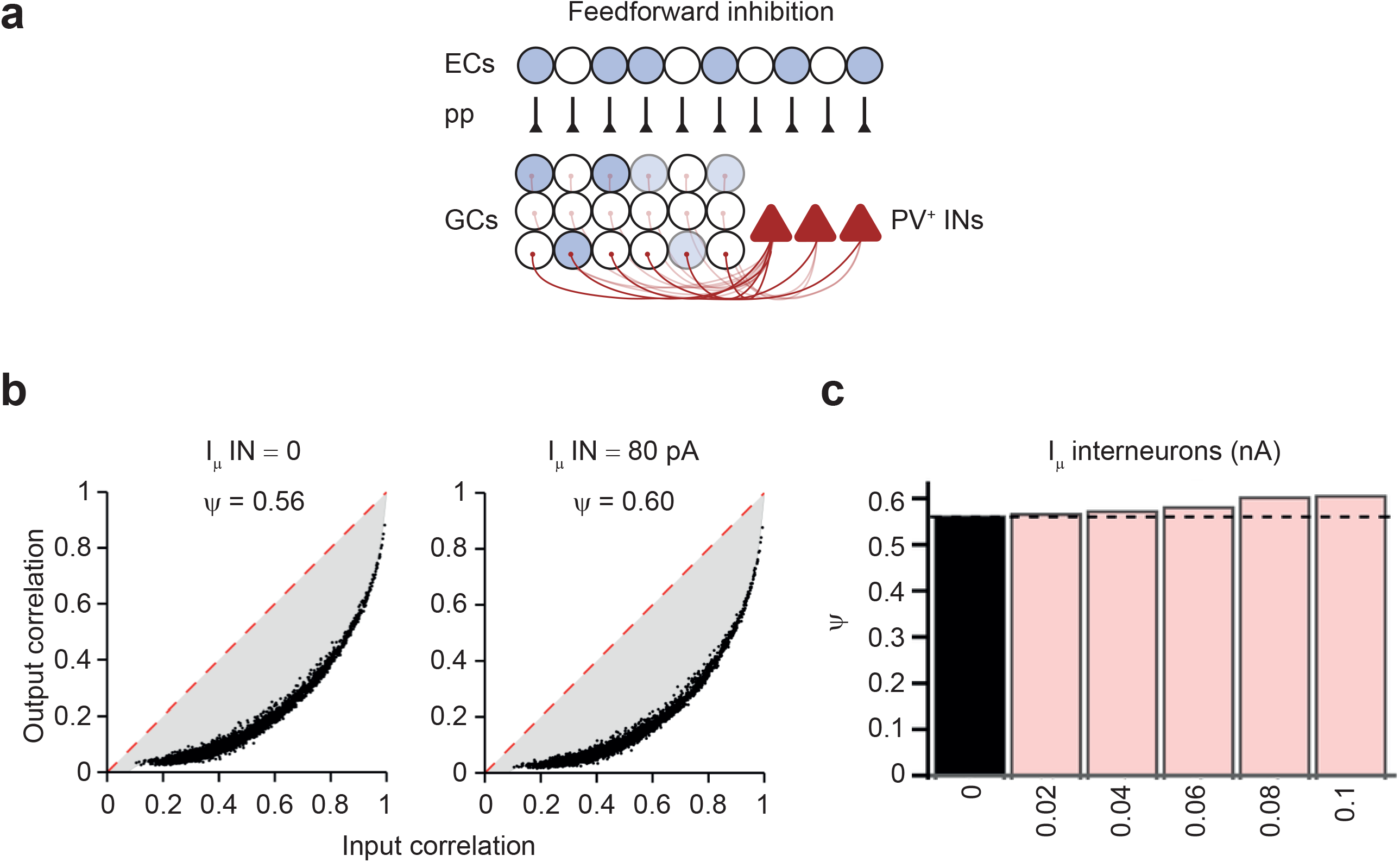
A network model incorporating feedforward inhibition is able to perform efficient pattern separation. (**a**) Schematic illustration of the network model incorporating feedforward inhibition, in addition to feedback inhibition. ECs innervate GCs and INs with similar connectivity rules. The tonic excitatory drive in an individual IN is computed from the drive from the nearest GC as:

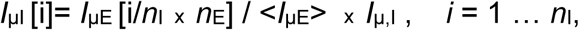

where *I*_µI_ [i] is the excitatory drive in the *i*^th^ interneuron (unitless), *I*_µE_ [i] is the excitatory drive in the i^th^ GC, *n*_I_is the number of INs, *n*_E_ is the number of GCs, <*I*_µE_> is the average excitatory drive over all GCs, and *I*_µ,I_ is the chosen excitatory drive in the INs (in pA). (**b**) Input-output correlation curves in a control network (left) and a network incorporating feedforward drive to INs (right). (**c**) Dependence of Ψ on feedforward drive in INs. Black bar, default value (no feedforward drive); light red bars, larger values (increased feedforward drive). Note that efficacy pattern separation Ψ is slightly increased by incorporation of feedforward excitation of INs.

**Supplementary Figure 4.**
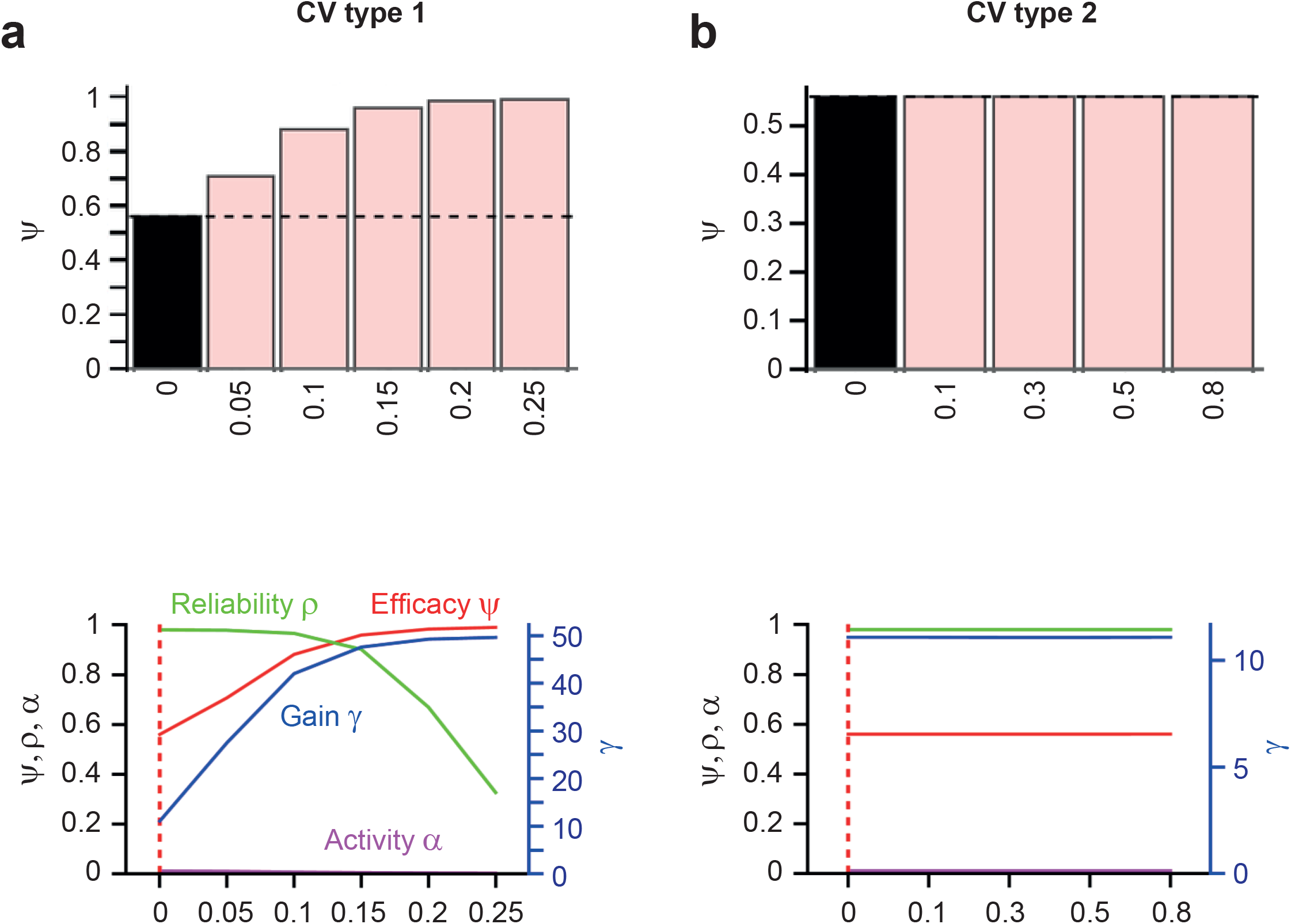
A network model incorporating type 1 and type 2 synaptic amplitude variability is able to perform efficient pattern separation. (**a**) Ψ for different degrees of type 1 (trial-to-trial) variability in the amplitude of all synapses (excitatory E–I, inhibitory I–E, and inhibitory I–I synapses). Coefficient of variation (CV = standard deviation / mean) was varied between 0.05 and 0.25. The synaptic weights fluctuated randomly from trial to trial. Top, summary bar graph of Ψ. Note that the type 1 variability resulted in an apparent increase in pattern separation efficacy Ψ, because an additional random process is added to the system. Bottom, plot of pattern separation efficiency Ψ (red), reliability ρ (green), gain γ (blue), and average activity α (magenta) as a function CV. Note that reliability ρ declines above a CV value of ~0.1. (**b**) Ψ for different degrees of type 2 (synapse-to-synapse) variability in the amplitude of all synapses (excitatory E–I, inhibitory I–E, and inhibitory I–I synapses). Coefficient of variation (standard deviation / mean) was varied between 0.1 and 0.8. The synaptic weights differed between individual synapses, but were constant from trial to trial. Note that type 2 variability has only minimal effects on pattern separation.

**Supplementary Figure 5.**
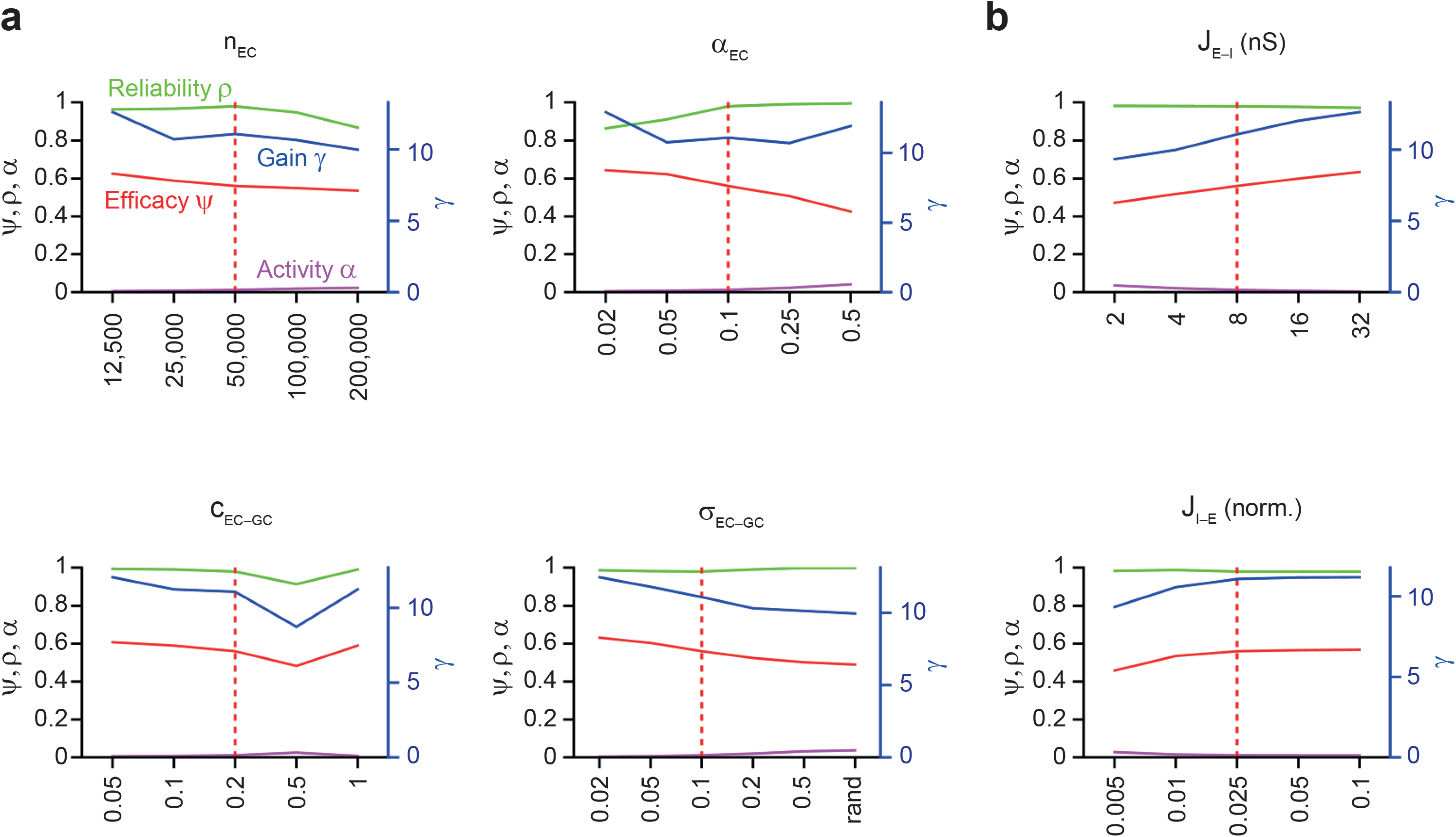
Efficiency, reliability, and gain of pattern separation after changes in EC–GC connectivity and synaptic strength of excitatory and inhibitory synapses. (**a**) Plot of pattern separation efficiency Ψ (red), reliability ρ (green), gain γ (blue), and average activity α (magenta) for different *n*_EC_, α_EC_, *c*_EC–GC_, and σ_EC–GC_ values (see Fig. 3c). (**b**) Plot of pattern separation efficiency Ψ (red), reliability ρ (green), gain γ (blue), and average activity α (magenta) for different *J*_E–I_ and *J*_I–E_ values (see Fig. 3d).

**Supplementary Figure 6.**
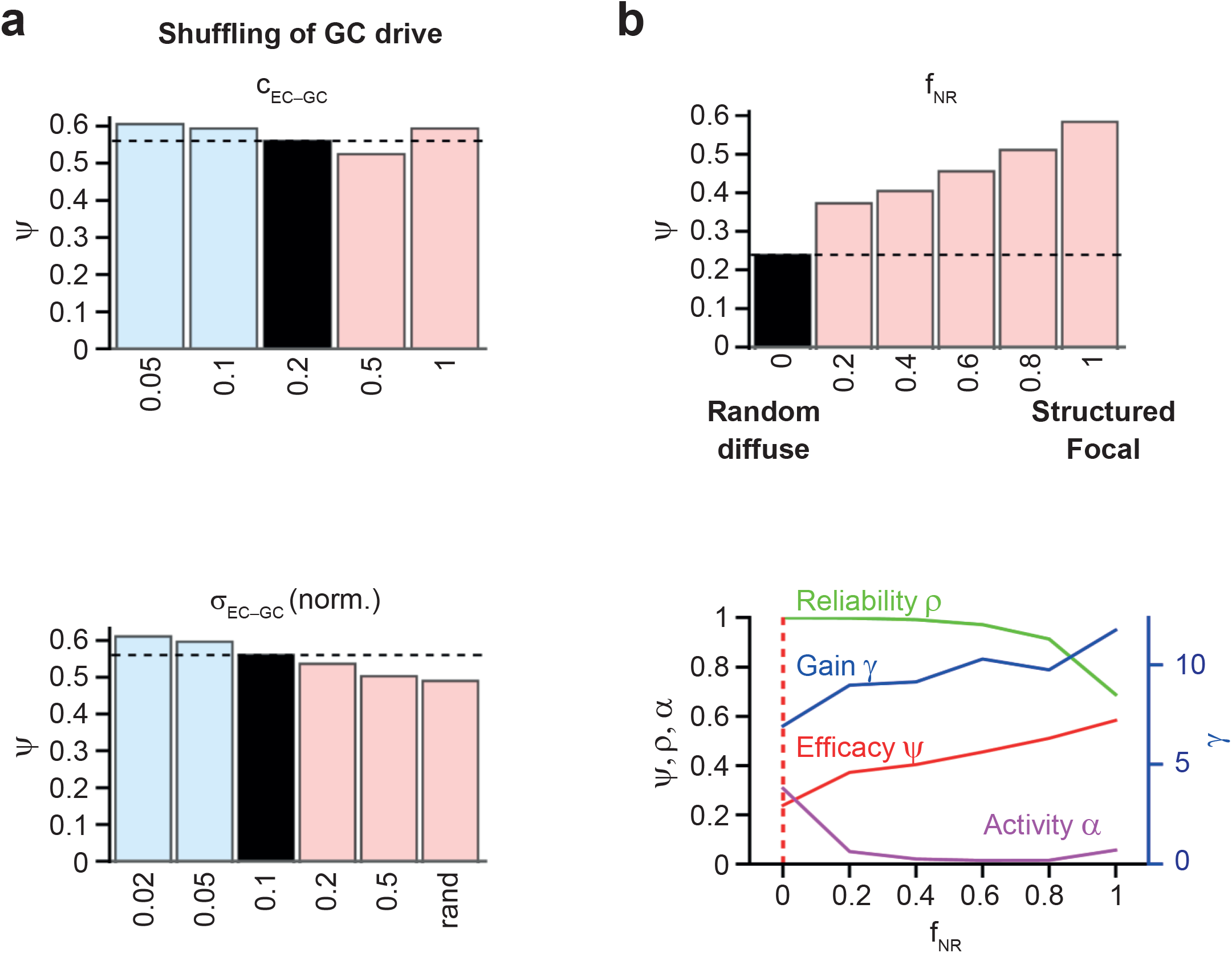
Effects of structured EC–GC connectivity on pattern separation. (**a**) Ψ for different values of peak connection probability (*c*_EC–GC_, top) and width of entorhinal connectivity (σ_EC–GC_, bottom) for a network in which the excitatory drive in GCs was randomly shuffled. Note that the results are only minimally different from those in the standard network (compare with Fig. 3c, bottom). (**b**) Effects of structural connectivity rules of EC–GC connections. EC–GC connectivity was either completely random, completely structured so that EC–GC synapses were formed within a full connectivity disc, or showed mixed properties. Top, bar graph of pattern separation efficiency Ψ for different proportions of nonrandom connections. Black bar, fraction of nonrandom connections *f*_NR_ = 0; i.e. all connections random; light red bars, enhanced structured connectivity (*f*_NR_ = 0.2 to 1, 1 = all connections structured). Bottom, plot of pattern separation efficiency Ψ (red), reliability ρ (green), gain γ (blue), and average activity α (magenta) as a function of the fraction of nonrandom connections.

**Supplementary Figure 7.**
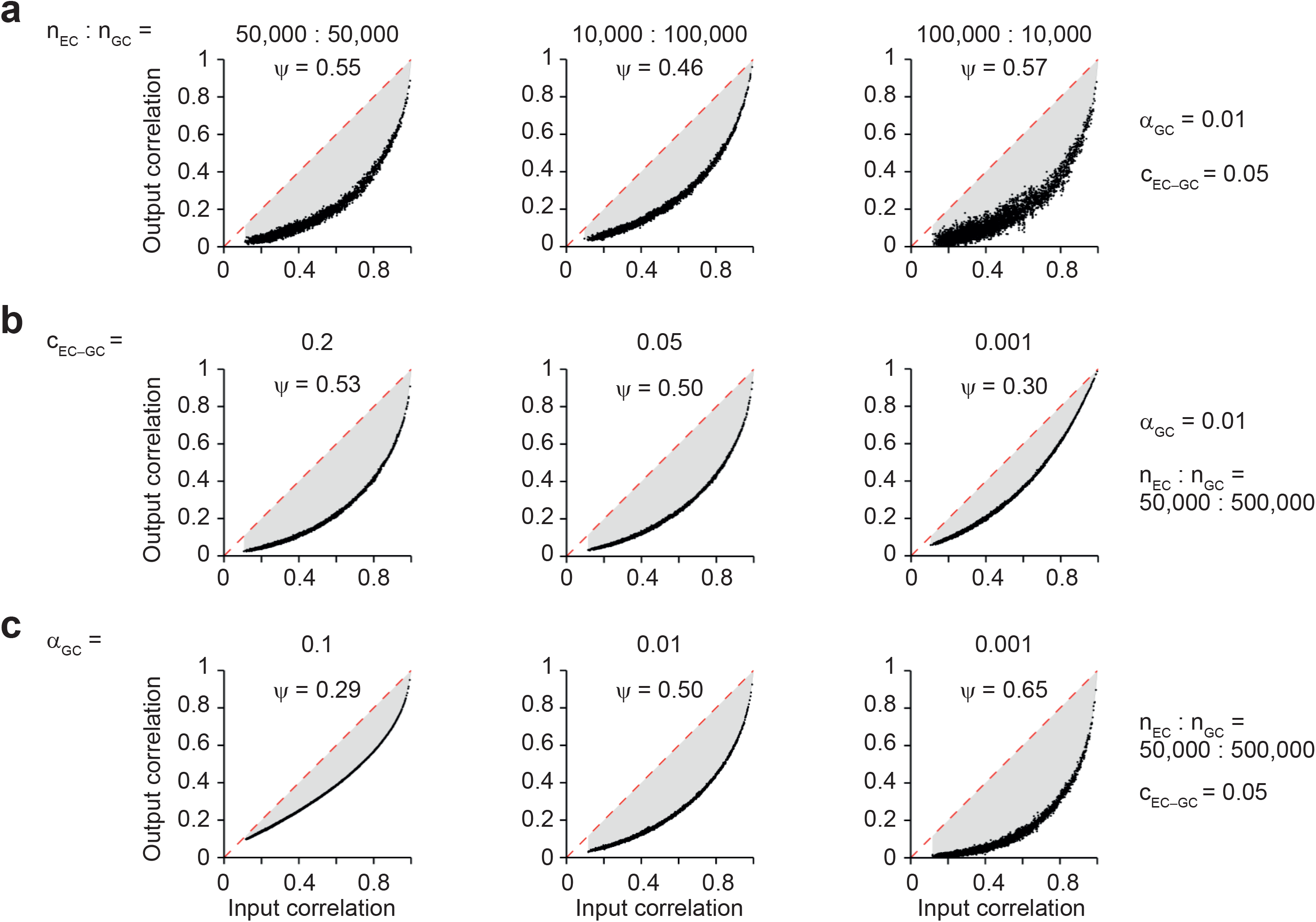
EC–GC connectivity rules have relatively minor effects on the efficacy of pattern separation. (**a**) Systematic analysis of effects of different expansion ratios on pattern separation. The ratio of cells in entorhinal and dentate gyrus layers, *n*_EC_: *n*_GC_, was varied over a wide range. Left, *n*_EC_: *n*_GC_ = 50,000: 50,000; center, *n*_EC_: *n*_GC_ = 10,000: 100,000; right, *n*_EC_: *n*_GC_ = 100,000: 10,000. Activity α_GC_ = 0.01, connection probability *c*_EC–GC_ = 0.05. Note that high Ψ values can be obtained at various expansion or compression ratios. (**b**) Systematic analysis of effects of EC–GC connection probability on the efficacy of pattern separation. Left, *c*_EC–GC_ = 0.2; center, *c*_EC–GC_ = 0.05; right, *c*_EC–GC_ = 0.001. Activity α_GC_ = 0.01, *n*_EC_: *n*_GC_ = 50,000: 500,000. Note that high Ψ values can be obtained for various values of EC–GC connection probability. (**c**) Systematic analysis of the GC activity level on the efficacy of pattern separation. Activity level α was set to 0.1, 0.01, and 0.001 (*n*_EC_: *n*_GC_ = 50,000: 500,000; *c*_EC–GC_ = 0.05). Activity in ECs was 0.1 in all simulations. Note that Ψ increased with decreasing α_GC_, i.e. increasing sparseness of activity in the dentate gyrus. The synaptic drive in GCs was computed as the product of entorhinal activity vector and EC–GC connectivity matrix, and thresholding was performed to obtain the desired average activity level in the GC layer.

**Supplementary Figure 8.**
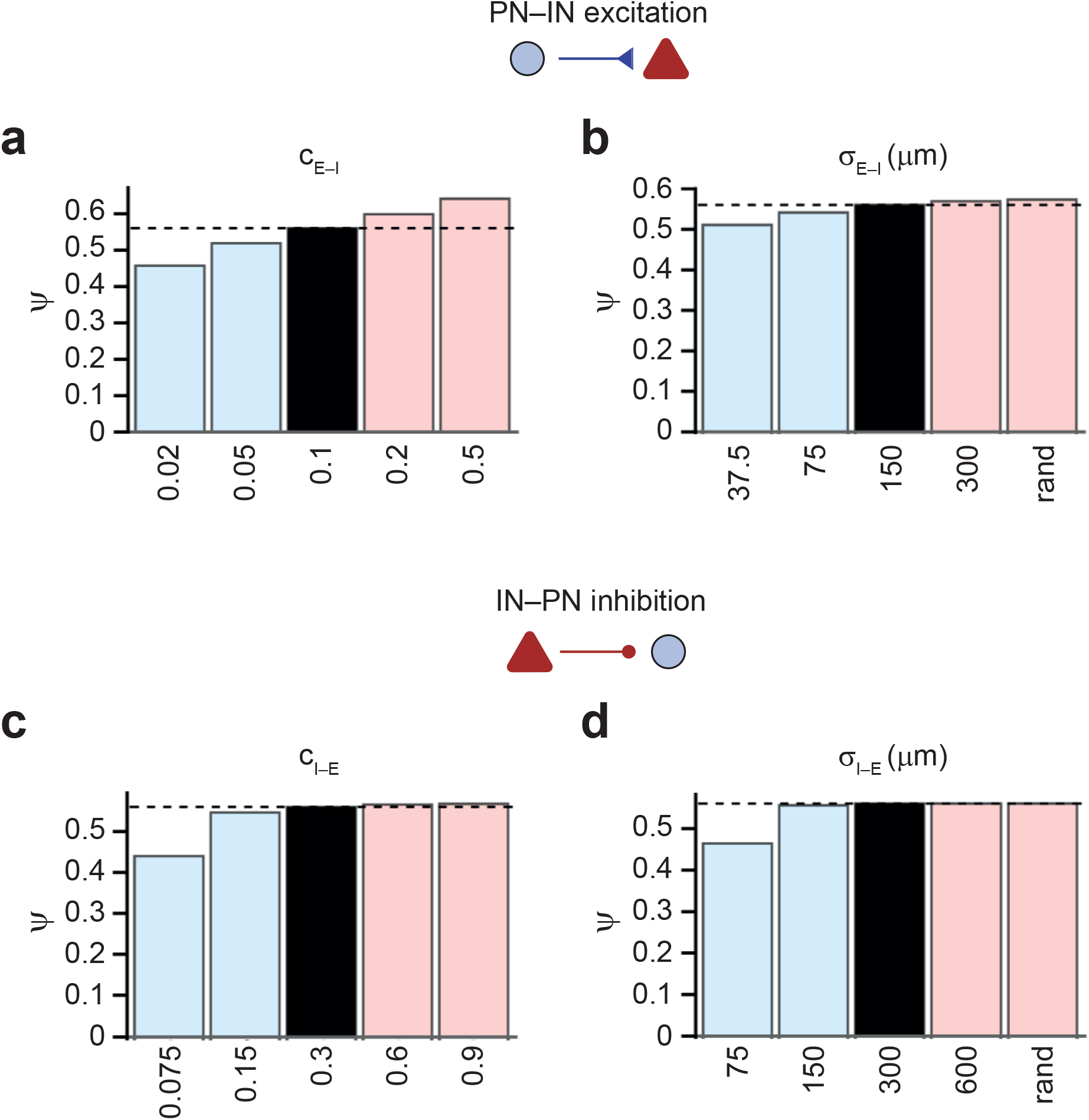
Interfering with lateral inhibition in different ways similarly affects pattern separation. (**a,b**) Ψ for different values of peak connection probability (*c*_E–I_, a) and width of excitatory E–I connectivity (σ_E–I_, b). (**c, d**) Ψ for different values of peak connection probability (*c*_I–E_, c) and width of inhibitory I–E connectivity (σ_I–E_, d). Note that the effects of changing excitatory and inhibitory connectivity on pattern separation efficacy are similar to those of changing the strength of excitatory E–I and inhibitory I–E synapses (Fig. 3d, right).

**Supplementary Figure 9.**
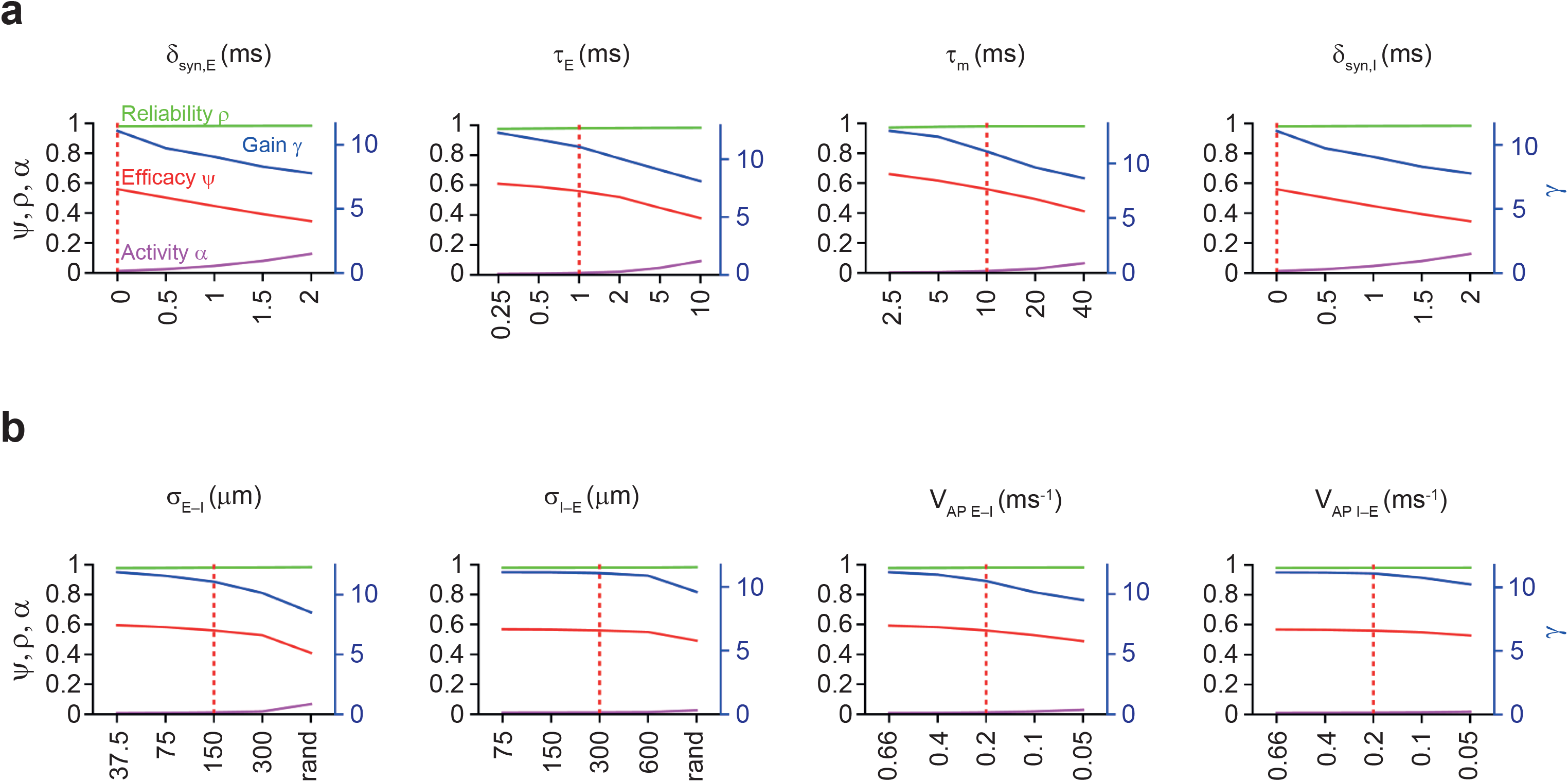
Efficiency, reliability, and gain of pattern separation after changes in fast signaling and focal connectivity of PV^+^ interneurons. (**a**) Plot of pattern separation efficiency Ψ (red), reliability ρ (green), gain γ (blue), and average activity α (magenta) for different δ_syn,E_, τ_decay,E_, τ_m_, and δ_syn,E_ values (see Fig. 4b). (**b**) Plot of pattern separation efficiency Ψ (red), reliability ρ (green), gain γ (blue), and average activity α (magenta) for different σ_E–I_, σ_I–E_, *v*_AP, E–I_, and *v*_AP, I–E_ values (see Fig. 4c and e).

